# Subspace-constrained approaches to low-rank fMRI acceleration

**DOI:** 10.1101/2020.12.15.422908

**Authors:** Harry T. Mason, Nadine N. Graedel, Karla L. Miller, Mark Chiew

## Abstract

Acceleration methods in fMRI aim to reconstruct high fidelity images from undersampled k-space, allowing fMRI datasets to achieve higher temporal resolution, reduced physiological noise aliasing, and increased statistical degrees of freedom. While low levels of acceleration are typically part of standard fMRI protocols through parallel imaging, there exists the potential for approaches that allow much greater acceleration. One such existing approach is k-t FASTER, which exploits the inherent low-rank nature of fMRI. In this paper, we present a reformulated version of k-t FASTER which includes additional L2 constraints within a low-rank framework.

We evaluated the effect of three different constraints against existing low-rank approaches to fMRI reconstruction: Tikhonov constraints, low-resolution priors, and temporal subspace smoothness. The different approaches are separately tested for robustness to undersampling and thermal noise levels, in both retrospectively and prospectively-undersampled finger-tapping task fMRI data. Reconstruction quality is evaluated by accurate reconstruction of low-rank subspaces and activation maps.

The use of L2 constraints were found to achieve consistently improved results, producing high fidelity reconstructions of statistical parameter maps at higher acceleration factors and lower SNR values than existing methods, but at a cost of longer computation time. In particular, the Tikhonov constraint proved very robust across all tested datasets, and the temporal subspace smoothness constraint provided the best reconstruction scores in the prospectively-undersampled dataset. These results demonstrate that regularized low-rank reconstruction of fMRI data can recover functional information at high acceleration factors without the use of any model-based spatial constraints.

Highlights

- We introduce an alternate implementation of low-rank fMRI reconstruction by using alternating minimization, which allows for easy integration of the subspace-specific L2 constraints
- We use the alternating minimization approach to accelerate FMRI by exploiting coil sensitivity, low-rank structures, and additional L2 constraints
- We found Tikhonov and Temporal Subspace Smoothness constraints show improved performance over other methods for R=15-30
- Tikhonov Constraints were the most robust of the constrained-subspace methods, with the shortest reconstruction time
- Temporal Subspace Smoothness produced the highest reconstruction scores in the prospectively under-sampled data

## 1. Introduction

fMRI is a non-invasive, whole-brain functional imaging technique that suffers from a trade-off between temporal and spatial resolution. Acceleration aims to increase the temporal resolution without loss of spatial resolution through higher sampling efficiency in conjunction with advanced image reconstruction that leverages additional information and/or constraints. By providing increased temporal degrees of freedom in a given scan duration, acceleration can: improve sensitivity to temporal features of the haemodynamic response; reduce physiological noise aliasing; and improve statistical power. Depending on the application, the increased sampling efficiency garnered from acceleration could also be used to reduce scan times, or to increase the spatial resolution.

Various acceleration techniques have been widely adopted for fMRI. Parallel imaging methods rely on the spatial variation of sensitivity profiles of multi-channel receiver coils, which provide additional spatial information in image reconstruction. This can occur in the image domain (e.g. SENSE [1]) or in the sampling domain (e.g. GRAPPA [2]). Simultaneous multi-slice imaging [3], [4] extends these in-plane techniques to accelerate across slices without reduction factor SNR penalties, increasing the achievable temporal resolution. Parallel imaging is conventionally a timepoint-by-timepoint approach that does not leverage any temporal information during reconstruction.

Methods which do jointly consider k-space and time are known as k-t methods and can be broadly separated into three categories: methods which make a strong assumption about the spatiotemporal structure [5]–[8], methods which make a strong assumption about sparsity within a pre-defined basis set (compressed sensing approaches) [9]–[13], and methods which assume the data fits a globally low-rank model [14], [15]. There are also approaches which combine these methods [13], [16]–[20]. By focusing on redundancies or structural features in k-t space, k-t methods have the potential for much greater degrees of acceleration than time-independent methods due to the extra dimension of shared information.

Compressed sensing approaches use L1 constraints methods to promote sparsity in reconstruction. These approaches have proven very effective in other fields of dynamic MRI reconstruction, but have had relatively limited adoption in fMRI, likely due to difficulty finding suitable sparse representations for the relatively subtle BOLD signals. While initial exploratory work in compressed sensing reconstruction for fMRI focused on spatial-domain sparse transformations [10], [11], most recent work incorporating sparsity assumptions have focused instead on sparsifying the temporal domain [21], [22]. Low rank + Sparse (L+S) methods [19], [20], are a recent set of combined approaches that aim to isolate the functional information in the sparse component of the reconstruction [23], [24] while capturing the non-sparse background in the low-rank component. The result of this approach is that the rank in the L component is kept very low and that the majority of the important BOLD information is in the S component, with PEAR [25] a notable recent example that explored the idea of capturing more BOLD information in the L component.

An alternative to sparse modelling of the BOLD signals is a conceptually simpler approach based on a regularized globally low-rank model of the fMRI data. There is a correspondence between the approaches that use training data to estimate a sparse or low-dimensional basis [13], [26] and low-rank models, since low-rank models by definition have few non-trivial components (i.e. the singular value distribution is sparse). However, low-rank models do not require prior knowledge of the sparse bases, and instead estimate the spatio-temporal basis representations for the data. The inherent low-rank nature of fMRI [15], which can be understood as the combination of a few spatially coherent temporal processes (i.e. activation maps that identify voxels with a common time-series), forms one such exploitable structure in a k-t representation of the data. In analysis of fMRI data, for example, a dimensionality reduction is often applied as a pre-processing step [27], which explicitly enforces a low-rank representation of the system prior to resting-state analysis methods such as independent component analysis (ICA) [28]–[30]. Various noise sources (e.g. thermal noise, physiological noise, etc.), motion, and image artefacts make the system only approximately low rank, although some confounds can also be estimated as low-rank processes [31].

Globally, low-rank methods can be used to represent space-time data as a spatial subspace paired with a temporal subspace and associated weighting factors. The Partially Separable Functions method (k-t PSF) [14], [32] is a data-driven approach that first identifies a temporal subspace from fully-sampled low spatial resolution and high temporal resolution training data, and then uses this to reconstruct a high resolution spatial subspace from under-sampled data. An alternative rank-constrained approach is k-t FASTER (fMRI Accelerated in Space-Time via Truncation of Effective Rank [15], [33]), which jointly identifies the subspaces that best describe the acquired data. Importantly, the only constraint imposed by k-t FASTER is that of fixed rank. The rank constraint alone is enough to achieve modest acceleration factors [15], but rank-constrained methods may also be combined with coil sensitivity information and non-Cartesian sampling [33] for increased acceleration.

In addition to the rank and coil sensitivity constraints, other information may also be incorporated into the reconstruction. Tikhonov regularization prevents overfitting on the temporal and spatial components, and serves as a way to penalize the energy content of the reconstruction. Radial k-space trajectories have a higher sampling density in central k-space than peripheral k-space, and so reweighting the low-resolution k-space could allow the reconstruction to be more strongly constrained in the densely sampled centre of k-space. The importance of central k-space more generally in MRI reconstruction has previously been used in approaches such as keyhole [8], k-t SPARSE [9], and k-t PCA [16]. Temporal regularization of some form has previously been incorporated into fMRI reconstruction in approaches like Dual TRACER [34] and temporal smoothness for simultaneous multi-slice EPI [35]. With a temporally varying sampling scheme, such as golden angle radial sampling (e.g. TURBINE [36]), enforcing temporal smoothness can be an effective way to reduce aliasing artefacts with a fractional penalty to the resulting temporal degrees of freedom.

In this paper, we explore extensions to the k-t FASTER approach that are formulated within an alternating minimization framework that incorporates L2-based regularization in addition to the previous fixed-rank constraints. We explore specific L2 constraints that correspond to Tikhonov regularization, low-resolution priors, and temporal subspace smoothness. Using L2-based constraints allows for interpretations of the constraints as Gaussian priors, and they are robust and relatively simple to implement. We compare the proposed approaches to unconstrained k-t FASTER and k-t PSF reconstructions of retrospectively and prospectively under-sampled datasets, which can be conceived of as special cases within this regularization framework. We evaluate these different methods with regards to the accuracy of the spatial and temporal components, and the sensitivity and specificity of statistical parameter maps (activation).

## 2. Material and methods

### 2.1 Theory

#### 2.1.1 Reformulation of k-t FASTER

The original k-t FASTER methodology used an iterative hard threshold + matrix shrinking approach [15] to enforce a fixed low-rank constraint on the reconstructed image time series. To enable us to easily introduce additional constraints on the spatial and temporal subspaces, we reformulate this low-rank optimization as a matrix factorization problem:

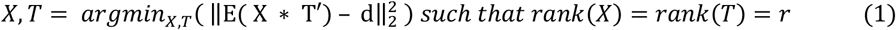

Equation 1 uses the following variables - E: sampling and multi-coil encoding function; d: multi-coil under-sampled k-t fMRI data; X: spatial components of decomposition; T: temporal components (T’ = Hermitian adjoint of T); ‖ ‖_2_: L2 norm, and r: rank constraint. For non-Cartesian sampling, E will contain an NUFFT operator [37]. The rank constraint will also apply to equations 2-5, but will be omitted for.

To solve the non-convex low-rank reconstruction, a minimization approach is used which alternately optimizes two convex subproblems [38]. These subproblems solve for either the spatial (X) or temporal (T) components, respectively, while the other variable is fixed. The spatial dimensions are vectorized, such that the product X*T’ forms a 2D space-time low-rank matrix that is our estimate of the fMRI time-series, and the 3D image volumes are a re-formatting of the 1D spatial vector. The decomposed matrices X and T form a low-rank decomposition, with the low-rank structure encoded in the dimensionality of the matrices, and X and T are not forced to be orthogonal. Pseudocode is included in Appendix A, and full implementation details are included in Appendix B.

#### 2.1.2 Soft Constrained-Subspace Approaches

The alternating minimization approach allows us to easily add additional subspace-specific constraints into Eq. 1, with the relative balance of low rank and additional constraints controlled by regularization parameters (λ). The original k-t FASTER approach can be reformulated by setting λ=0 in all the following equations. Formulations with non-zero and non-infinity λ will be referred to as softly constrained. Figure 1 contains schematics which demonstrate the various approaches.

**Figure 1:**
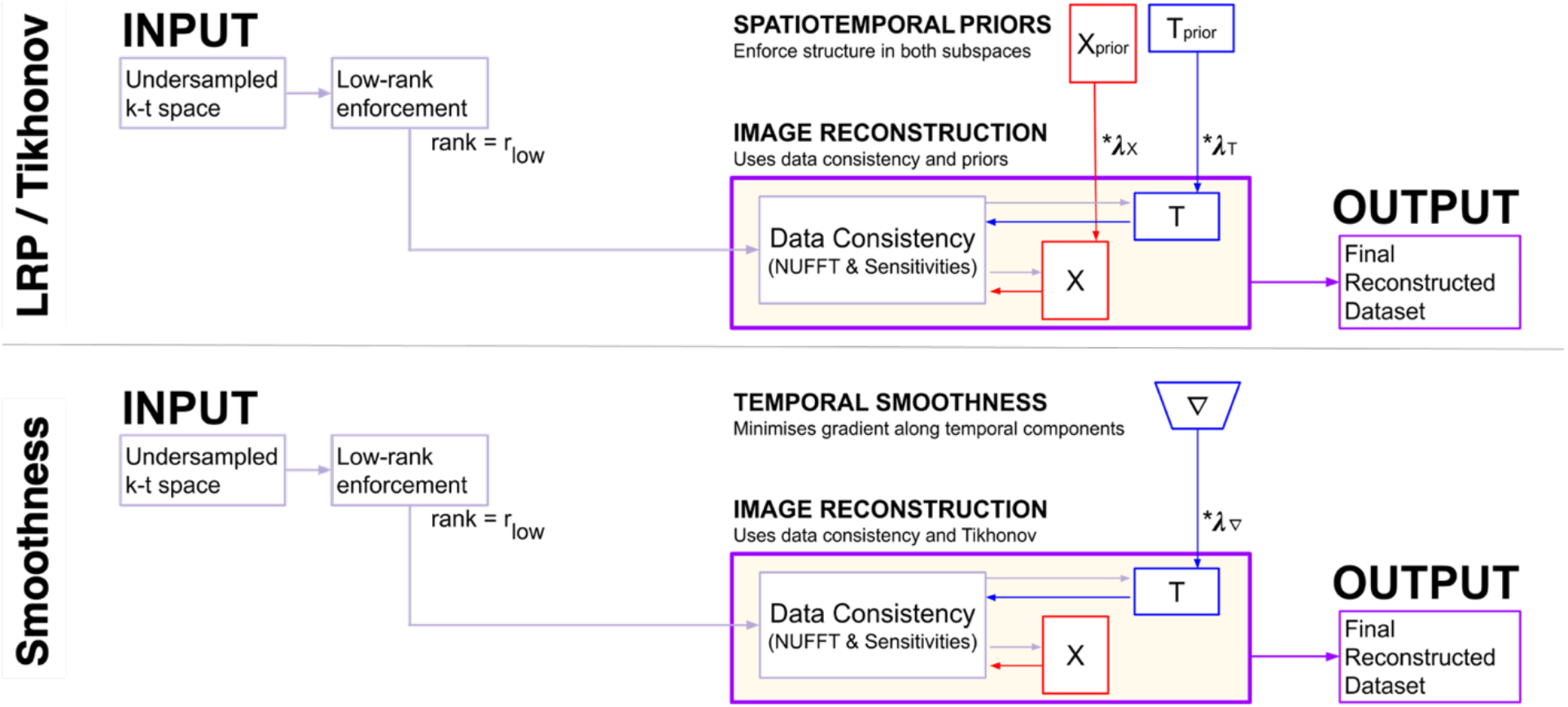
A schematic overview of a reconstruction with various constrained-subspace approaches. For the LRP, X_prior_ and T_prior_ are created using a windowed version of the under-sampled data according to only the rank constraints and coil sensitivity information. For Tikhonov, X_prior_ and T_prior_ are zero-filled. X_prior_ and T_prior_ are fed as a constraint into the final reconstruction, combining with the data consistency term on an unwindowed dataset to produce the final output. The temporal subspace smoothness schematic shows a finite difference matrix ∇ applied solely to the temporal component matrix T, before also being combined with the data consistency term.

##### Tikhonov

The most straightforward constrained-subspace approach derives from methods used for collaborative filtering [39], which often uses Tikhonov regularization on the two component matrices (X and T). L2-regularization terms are included to serve as energy minimization terms for each variable, which prevent matrix entries from becoming too large:

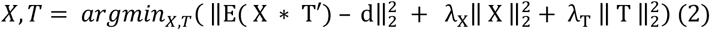

##### Low-Resolution Priors

For data acquired using trajectories with non-uniform sampling densities that sample the centre of k-space each TR, one can formulate a L2 regularization corresponding to Low-Resolution Priors (LRP). In uniform radial sampling drawn from multiple spokes (TRs) within a plane, a central window of radius 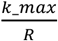 fulfils the Nyquist sampling criteria in the azimuthal direction. Additionally, these low spatial frequencies represent the net balance of temporal processes at the ultimate temporal resolution, but without capturing detailed spatial features. This central window can be more strongly weighted during a final reconstruction to accurately capture these high temporal resolution processes.

The LRP constraints (X_prior_ and T_prior_) are created by windowing the full k-space dataset with a Tukey window 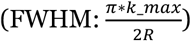 and then reconstructing X and T using Equation 1, albeit with *d* referring to windowed k-space data, analogous to the estimation of the temporal subspace from training data in the k-t PSF approach. The final reconstruction is then weighted by the LRPs along with the full unwindowed sampled data (Equation 3).

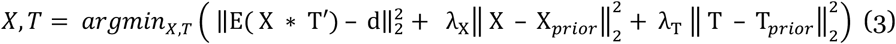

The previously proposed k-t PSF method represents a special case of the more general LRP framework. This method reconstructs the spatial coefficients against a temporal basis (or prior) estimated from low-resolution training data. k-t PSF can be formulated in the Eq. 3 framework by setting λ_X=0_ and λ_T =∞_. The temporal subspace is constrained to be identical to this predetermined basis, which is labelled T_prior_:

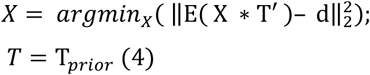

##### Temporal Subspace Smoothness

The aim of a temporal subspace smoothness term is to preserve the relatively smooth BOLD response (particularly at high acceleration) and reduce the magnitude of high temporal frequency under-sampling artefacts. Trajectories with a sampling point-spread function that changes every frame (e.g. golden angle radial trajectories) can result in high temporal frequency under-sampling artefacts, and so are well suited to this approach. The reconstruction is governed by equation 5. ∇ is a finite difference operator acting on the temporal dimension of each temporal process, and λ_∇_ is the corresponding weighting parameter:

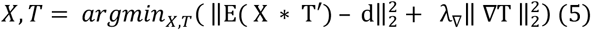

### 2.2 Experimental Details

We evaluated the different reconstructions (Tikhonov-constrained, LRP-constrained, smoothness-constrained, k-t FASTER, and k-t PSF) with both retrospectively under-sampled data in various SNR regimes, and with prospectively under-sampled data. The reconstructions are evaluated based on how accurately the spatial, temporal, and functional information is captured across a range of acceleration factors.

#### 2.2.1 Data Acquisition

In order to fulfil the non-uniform sampling requirements of the LRP constraints and the changing sampling PSF requirement of the smoothness constraints, all acquisitions in k-space followed the TURBINE trajectory [36], [40], a 3D hybrid radial-Cartesian EPI sequence which rotates an EPI blade around the phase encoding axis at constant azimuthal increments of the Golden Ratio angle (π/Φ ≈ 111.25°) [41]. This scheme provides a near-uniform radial sampling of k-space from any arbitrary post-hoc combination of consecutive blades, allowing for flexible degrees of acceleration (Figure 2) [42]. The under-sampling (or acceleration) factor R is defined here as the ratio of sampling lines required to fully sample k-space to the number of sampling lines acquired. In radial sampling, R=1 requires π/2 times more lines than Cartesian sampling.

**Figure 2:**
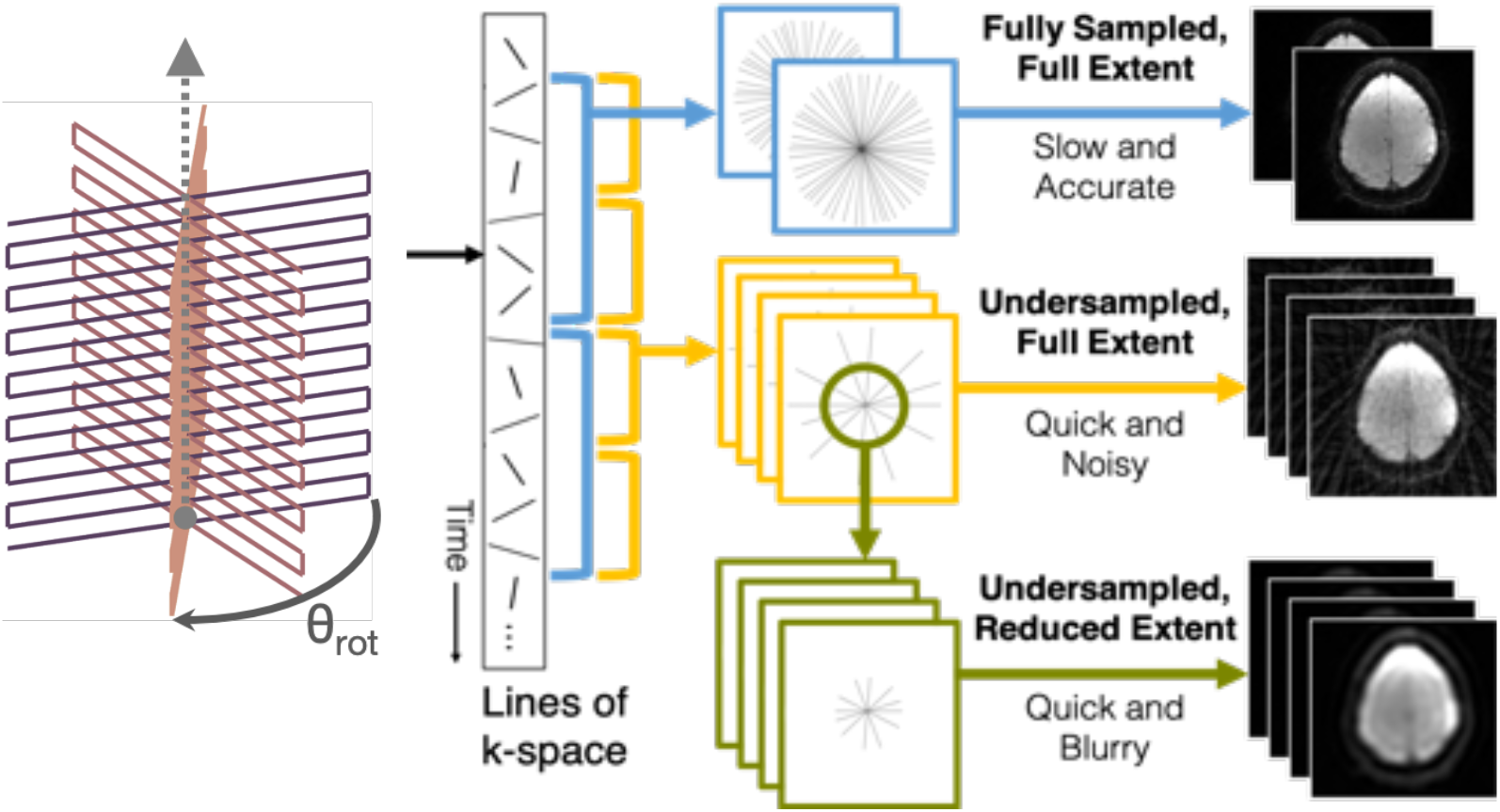
A demonstration of the flexibility of a golden angle sampling scheme, and of the k-space windowing required to create LRP constraints. EPI planes (left) is rotated by ≈ 111.25° around the phase-encoding axis. These rotated planes can then be flexibly combined. If many planes are used (top, blue) then a clean image is easily generated, but at the cost of temporal resolution. If fewer planes are used (middle, yellow) then more images are generated per second, but with an increased number of artefacts. The central part of under-sampled k-space satisfies the Nyquist criterion, even if the full extent of the under-sampled k-space does not. By windowing this central k-space (green, bottom), an accurate low-resolution depiction of the underlying data can be created.

All datasets were generated from a 30s/30s on/off finger-tapping task, and recreated 100×100 images with a 2mm isotropic voxel resolution. An SVD compressed the 32-coil channel to the 8 most dominant components for speed/memory purposes [43], [44]. All data were acquired on a 3T system (Prisma, Siemens Healthineers, Erlangen Germany) with informed consent in accordance with local ethics.

##### Retrospectively Under-sampled Datasets

“Retrospective dataset A” was created by retrospectively resampling each frame of a fully sampled dataset (300 frames, TR_frame_=1s) in k-space with a TURBINE pattern. The original dataset is used as a comparative ground truth, and was acquired as a full volume through a TURBINE acquisition with 20 blades/frame (TR_blade_=50ms, TE_blade_=30ms), and a single axial slice with clear bilateral activation was chosen for reconstruction. No rank reduction was applied to the original data. The dataset was sampled from a magnitude-only ground truth, with no added noise or phase variation. The retrospective acceleration factors used are R=15.71, 31.42, 39.27, and 52.36 (corresponding to 10, 5, 4, and 3 blades/frame respectively).

“Retrospective dataset B” was created by adding complex Gaussian noise in k-t space to retrospective dataset A at R=31.42, to highlight the performance difference between the different approaches with additional thermal noise. Noise was added to form new noisy datasets with high (SNR=100), medium (50) and low (20) SNRs, with the original dataset considered noiseless for the purposes of comparison. For each SNR, five unique instantiations of the noise were added to the underlying data before reconstruction. These values are representative of actual fMRI SNR values [45]. This additional Gaussian noise only models additive thermal noise as a step towards more realistic data (coherent noise sources such as physiological noise with temporal autocorrelation are not modelled here).

##### Prospectively Under-Sampled Data

The prospectively accelerated reconstructions used a TURBINE acquisition across eight different slices centred on the motor cortex. Slices were first reconstructed by performing an inverse FFT along the phase-encode (z) direction before a k-t reconstruction was carried out on each (x-y) k-space plane. Identical acquisition parameters with the same experimental set-up (TR_blade_=50ms, TE=30ms, flip angle=15°, BW=1786 Hz/px) were used for a short experiment (320s, five 30s on/off task epochs) and a long experiment (640s, ten epochs) which were carried out consecutively on the same subject. An R=1.05 reconstruction of the long dataset contains enough temporal Degrees-of-Freedom to characterize the underlying functional signal and provide high-quality activation maps, serving as a fully-sampled approximate "ground truth" reference against which the reconstruction of the accelerated short dataset is compared. The different acceleration factors in the short prospective dataset (R=7.85, R=15.71, R=26.18) lead to different temporal resolutions and temporal degrees of freedom, as well as affecting other statistical properties (such as physiological noise variance). While the most general method would reconstruct all eight slices simultaneously to capture shared temporal processes, the extra computational power required for this was not considered worth the benefits, and hence slices were reconstructed independently. The reconstruction details are listed in table 1.

**Table 1:** The reconstruction details for the different acceleration factors used in reconstructing the prospectively under-sampled data.

| DATASET | BLADES | $TR_{FRAME}(S)$ | R | BLADES/FRAME | FRAMES |
| --- | --- | --- | --- | --- | --- |
| LONG | 12,800 | 7.5 | 1.05 | 150 | 85 |
| SHORT | 6,400 | 1 | 7.85 | 20 | 320 |
| SHORT | 6,400 | 0.5 | 15.71 | 10 | 640 |
| SHORT | 6,400 | 0.3 | 26.18 | 6 | 1066 |

#### 2.2.2 Selection of reconstruction parameters

A logarithmic grid search over potential λ_X_ and λ_T_ candidates was carried out for all datasets, constraints, and acceleration factors. The grid search for retrospective dataset A is shown in Figure 3 to demonstrate the typical effects of varying λ on the reconstructed spatial and temporal information for the different constraints, with boundary cases shown for λ=0 (zero prior influence) and λ=∞ (the solution is fixed to the prior). The special boundary case of (λ_X_ = 0, λ_T_ = 0) defines k-t FASTER for all constraints and the special case of (λ_X_ = 0, λ_T_ = ∞) defines k-t PSF with LRP constraints. As the smoothness constraints rely on a single weighting parameter (λ_∇_), the results are shown as a line graph.

**Figure 3:**
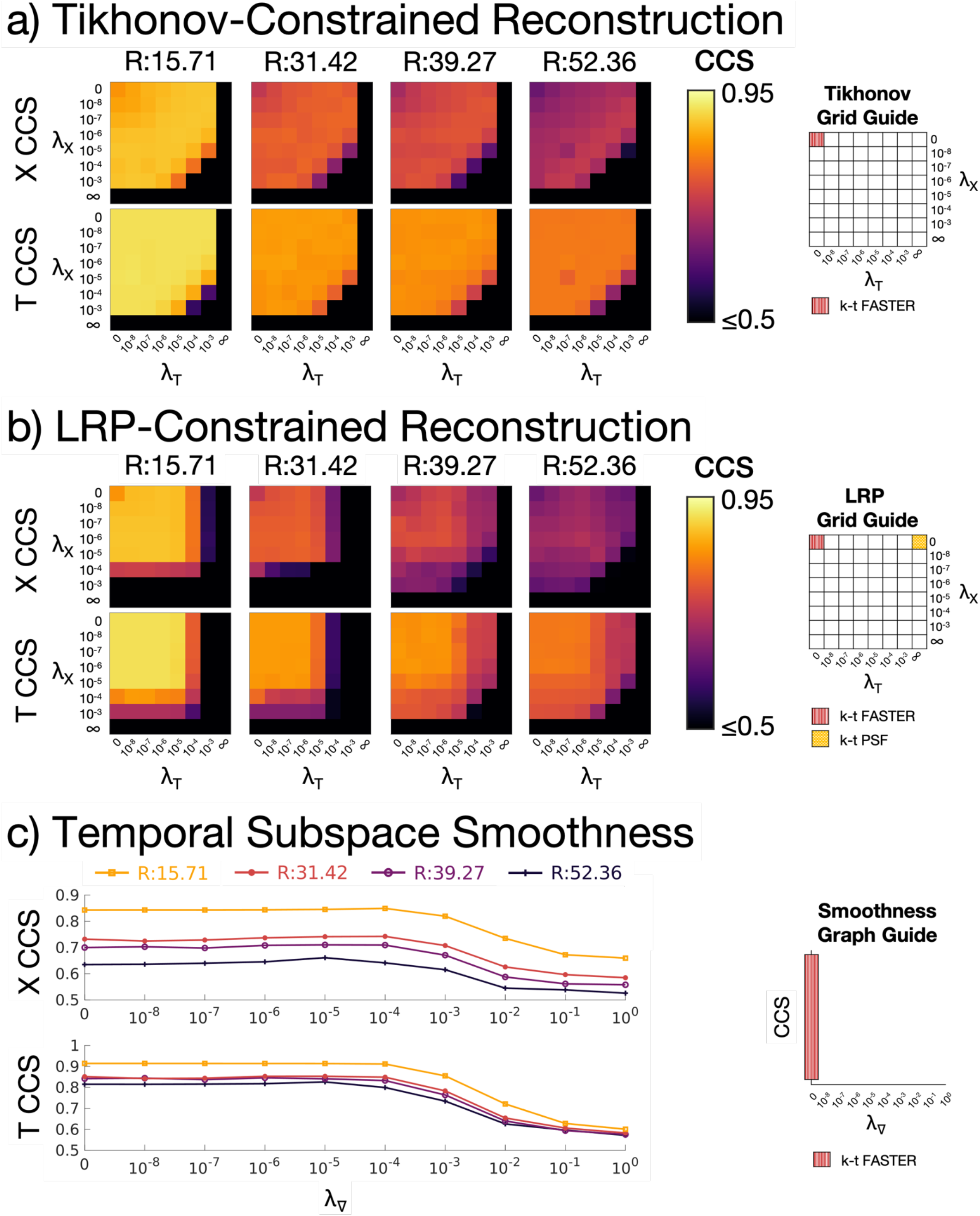
The canonical correlation scores (CCS) of retrospective dataset A vs a ground truth for a): Tikhonov-constrained reconstructions, b): LRP-constrained reconstructions, c): Temporal Subspace Smoothness reconstructions. X CCS and T CCS refer to the spatial and temporal Canonical Correlation Scores respectively. The acceleration factors shown are: R=15.71 (10 blades/frame), R=31.42 (5 blades/frame), R=39.27 (4 blades/frame), and R=52.36 (3 blades/frame). The λ values encoding the pre-existing k-t FASTER and k-t PSF methods are shown on the right for each constraint.

The reconstruction rank was fixed at 16 in all cases (a value used in recent literature for low-rank task fMRI [46]), and a variety of acceleration factors were tested. The convergence criterion was defined as the normalized gradient for the whole cost function *CF* (equation 6), evaluated after the temporal subproblem optimization for iteration number *i*.

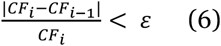

A criterion of ε = 10^−5^ was used for both retrospective datasets, which was chosen as the value at which a k-t FASTER reconstruction with different random initializations was found to converge to identical subspaces. For the prospective dataset, ε = 10^−3^ was found to be more optimal. This lower convergence criterion was found to produce slightly improved statistical parameter maps (defined using the metrics of section 2.2.3), which may be a result of overfitting occurring at the more precise criterion used in both retrospective datasets. The different criteria chosen here were selected to ensure a very high level of agreement regardless of the initialization, and was chosen using the k-t FASTER reconstruction without additional subspace constraints. Future experiments may well benefit from more liberal criteria to enable faster reconstruction, without necessarily experiencing any loss in reconstruction quality.

#### 2.2.3 Evaluation and fMRI Analysis

Reconstruction image quality can be difficult to determine [47], with more incoherent (‘noise-like’) artefacts usually preferable to coherent artefacts, and the first component of the subspace dominating most image quality metrics (such as root mean square error or structural similarity index). Spatial artefacts can also make conventional metrics like SNR (or simple measures of noise) harder to quantify.

Instead, the spatial and temporal subspaces were directly compared to the retrospective ground truth subspaces using canonical correlation analysis. Canonical correlation measures the cosine of the principal angles (the alignment) between subspaces [48], with higher values reflecting more aligned subspaces, and a value equal to the rank of the subspace (16 in all cases) demonstrating complete alignment. A Canonical Correlation Score (CCS) was created by dividing the canonical correlation by the maximal rank of the decomposed matrices, providing a normalized metric measuring the alignment of the subspaces. X CCS and T CCS respectively refer to the CCS for spatial and temporal subspace analyses. As a subspace alignment metric, CCS does not account for the magnitude of the estimated components, only their relative alignment. This potential shortcoming is accepted for two reasons: firstly the data consistency term will generally ensure that the relative magnitude of the signal is well captured, and secondly any ICA analysis run on the data will also be scale-independent [49].

For all datasets, task fMRI analysis was performed in FEAT (FSL) [50]. To account for residual autocorrelation, the resulting z-statistic maps were null-corrected using mixture modelling [29], and the reconstructed prospective data is aligned to the ground truth reference using FLIRT [51] prior to analysis. Receiver Operating Characteristic (ROC) curves were calculated to measure the false positive rate (FPR) against true positive rate when comparing the reconstructions against the activation map of a fully sampled reconstruction. A threshold of z>3.1 was used to threshold the retrospective truth, and z>4.8 was used for the prospective data (these values were selected heuristically based on anatomical veracity of known regions of expected activation). Z-statistic parameter maps are shown at a false positive rate of 0.0015 in order to facilitate visualization. The ROC curves will be focussed on low FPRs, as the z-statistic corresponding to high FPRs would never be used in studies. The Area Under the Curve (AUC) of the full ROC curve allows for a simple comparison of many reconstructions, but the underlying z-statistic maps also provide valuable information as to the spatial location of false positives and false negatives.

## 3. Results

Optimal values of λ_X_, λ_T_, and λ_∇_ are evaluated for each dataset, method, and acceleration factor, and then the optimized reconstructions are evaluated against the reconstructions using the k-t FASTER and k-t PSF methods. The optima are selected using a heuristic combination of the CCSs, ROC AUCs, and qualitative assessments of z-statistic activation maps.

### 3.1. Retrospective Dataset A Results

The influence of λ_X_ and λ_T_ on the recovered temporal and spatial components for different constraints is shown in Figure 3. The LRP constraints are defined by a peak in spatial CCS and a broad plateau in temporal CCS (although the gradient is quite shallow near the peak). The Tikhonov constraints were defined by a line of peak values normal to λ_X_ = λ_T_, suggesting a 1D search could suffice to find an optimal λ pairing. For Tikhonov and LRP constraints, the upper-left-hand corner of every λ grid represents k-t FASTER, and the far left point represents k-t FASTER in the 1D plot. The upper-right-hand corner of the LRP constraint λ grids represent k-t PSF. The optimal λ values are shown in table 2, and were constant across acceleration factors, except for the highest acceleration factor (R=52.36).

**Table 2:** The optimal λ values for each method in retrospective dataset A. Results within 0.001 of the best ROC AUC score and 0.01 of the best CCS values are shown in bold.

| R | Method | $\lambda_x$ | $\lambda_T$ | $\lambda_v$ | X CCS | T CCS | ROC AUC |
| --- | --- | --- | --- | --- | --- | --- | --- |
| 15.71 | Tikhonov | $10^{-5}$ | $10^{-5}$ | 0 | **0.89** | **0.91** | **0.9983** |
| | LRP | $10^{-5}$ | $10^{-5}$ | 0 | **0.88** | **0.91** | **0.9985** |
| | Smoothness | 0 | 0 | $10^{-5}$ | 0.85 | **0.91** | **0.9983** |
|  | k-t FASTER | 0 | 0 | 0 | 0.84 | **0.91** | **0.9983** |
| | k-t PSF | 0 | ∞ | 0 | 0.34 | 0.28 | 0.8956 |
| 31.42 | Tikhonov | $10^{-5}$ | $10^{-5}$ | 0 | **0.80** | **0.85** | **0.9984** |
| | LRP | $10^{-5}$ | $10^{-5}$ | 0 | 0.78 | 0.82 | **0.9984** |
| | Smoothness | 0 | 0 | $10^{-5}$ | 0.74 | **0.85** | **0.9975** |
|  | k-t FASTER | 0 | 0 | 0 | 0.73 | **0.85** | 0.9973 |
| | k-t PSF | 0 | ∞ | 0 | 0.22 | 0.20 | 0.7052 |
| **39.27** | Tikhonov | $10^{-5}$ | $10^{-5}$ | 0 | **0.76** | **0.83** | **0.9974** |
| | LRP | $10^{-5}$ | $10^{-5}$ | 0 | 0.72 | 0.78 | **0.9968** |
| | Smoothness | 0 | 0 | $10^{-5}$ | 0.71 | **0.84** | 0.9956 |
|  | k-t FASTER | 0 | 0 | 0 | 0.70 | **0.84** | 0.9956 |
| | k-t PSF | 0 | ∞ | 0 | 0.21 | 0.22 | 0.5213 |
| **52.36** | Tikhonov | $10^{-5}$ | $10^{-5}$ | 0 | **0.73** | **0.82** | **0.9967** |
| | LRP | $10^{-4}$ | $10^{-6}$ | 0 | 0.67 | 0.78 | **0.9962** |
| | Smoothness | 0 | 0 | $10^{-4}$ | 0.65 | 0.80 | 0.9938 |
|  | k-t FASTER | 0 | 0 | 0 | 0.63 | **0.81** | 0.9927 |
| | k-t PSF | 0 | ∞ | 0 | 0.21 | 0.23 | 0.5741 |

Z-statistic activation maps were derived for all approaches using the optimized λ values at R=31.42 (Figure 4) and R = 52.36 (Figure 5). The ROC curves and activation maps are consistent with the results of Figure 3, with the Tikhonov and LRP constraints performing better than the other k-t methods at both acceleration factors, albeit with the Tikhonov regularization marginally outperforming LRP-constrained reconstruction at R=52.6. The cleanness of the dataset appeared to allow very high reconstruction factors which were not found to be possible in more realistic data.

**Figure 4:**
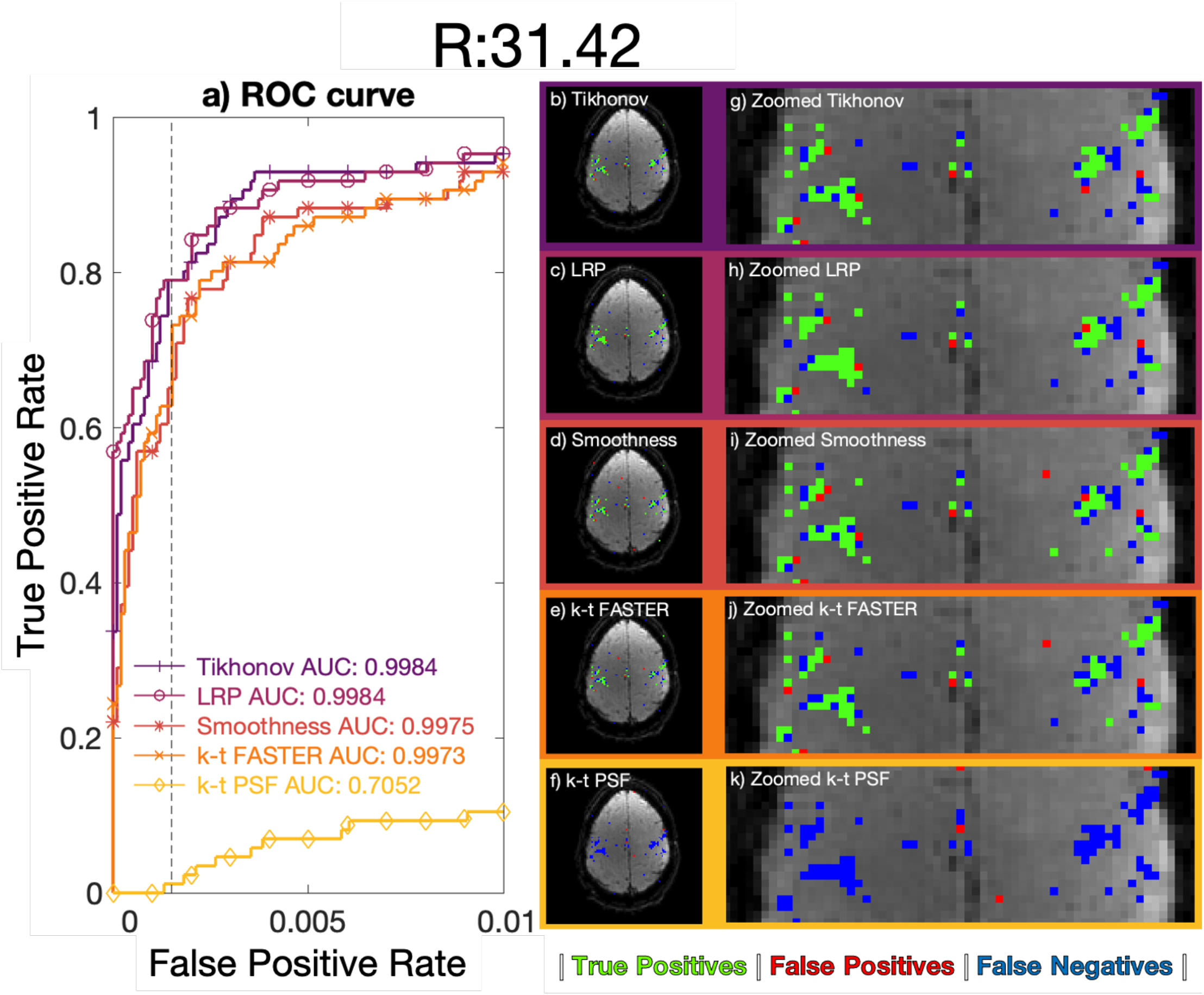
R=31.42 (5 blades/frame) retrospective dataset A reconstructions. a) ROC curves, legend lists full curve AUC. b)-f) Activation maps using a z-statistic corresponding to an FPR of 0.15%. g)-k) A medial zoom of the associated activation maps. b/g) Tikhonov: λ_X_ = 10^−5^, λ_T_ = 10^−5^, c/h) LRP: λ_X_ = 10^−5^, λ_T_ = 10^−5^, d/i) Temporal subspace smoothness: λ_∇_ = 10^−5^, e/j) k-t FASTER, f/k) k-t PSF. Maps b)-k) use green true positive pixels, red false positives, and blue false negatives.

**Figure 5:**
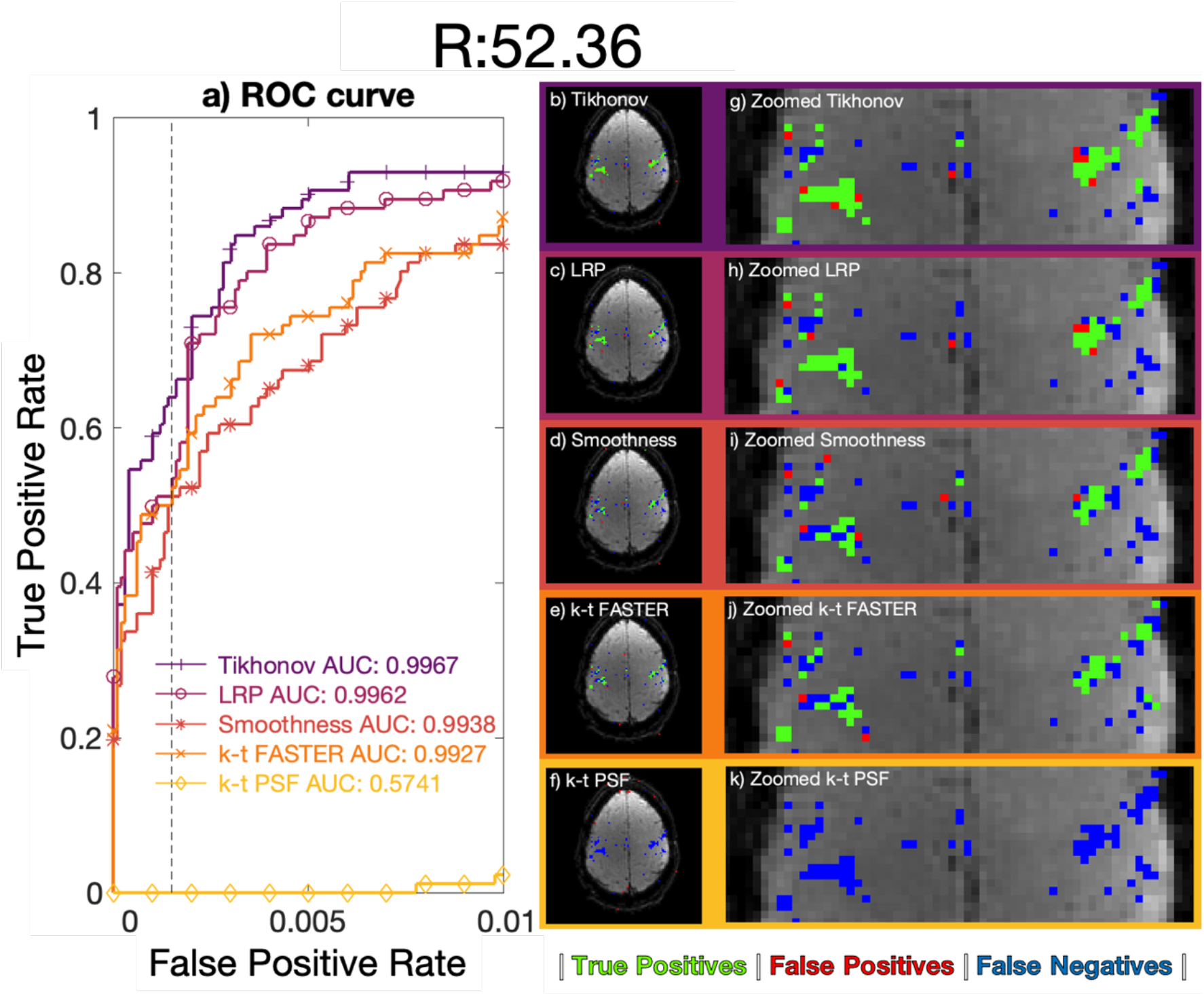
R=52.6 (3 blades/frame) retrospective dataset A reconstructions. a) ROC curves, legend lists full curve AUC. b)-f) Activation maps using a z-statistic corresponding to an FPR of 0.15%. g)-k) A medial zoom of the associated activation maps. b/g) Tikhonov: λ_X_ = 10^−5^, λ_T_ = 10^−5^, c/h) LRP: λ_X_ = 10^−4^, λ_T_ = 10^−6^, d/i) Temporal subspace smoothness: λ_∇_ = 10^−4^, e/j) k-t FASTER, f/k) k-t PSF. Maps b)-k) use green true positive pixels, red false positives, and blue false negatives.

### 3.2 Retrospective Dataset B Results

Optimal λ was found to increase as SNR decreased for Tikhonov and LRP results. The following values were used for both Tikhonov and LRP constraints: high SNR (SNR=100, λ_X_=10^−4^, λ_T_ =10^−5^); medium SNR (SNR=50, λ_X_=10^−4^, λ_T_ =10^−4^); low SNR (SNR=20, λ_X_=10^−3^, λ_T_ =10^−4^). The temporal subspace smoothness results used λ_∇_ = 10^−4^ in all cases, although the variation in results was small for 10^−4^ < λ_∇_ < 10^−1^.

The mean AUC of the noisy parameter map ROCs compared to a noiseless truth are summarized in Figure 6, with all reconstructions losing fidelity as SNR decreased. The noiseless reconstructions are equivalent to the data shown in figure 4. Maps comparing thresholded z-stat maps with the ground truth for each method are shown in Figure 7, with full visualizations of all reconstruction activation maps and ROC curves shown in Supplementary Figures 4-6. In t-tests performed between the different constraints within the three non-noiseless SNRs, all reconstructions within an acceleration factor were significantly different (p<0.05) except Tikhonov vs LRP at high SNR, k-t FASTER vs k-t PSF at low SNR, and LRP vs Smoothness at low SNR.

**Figure 6:**
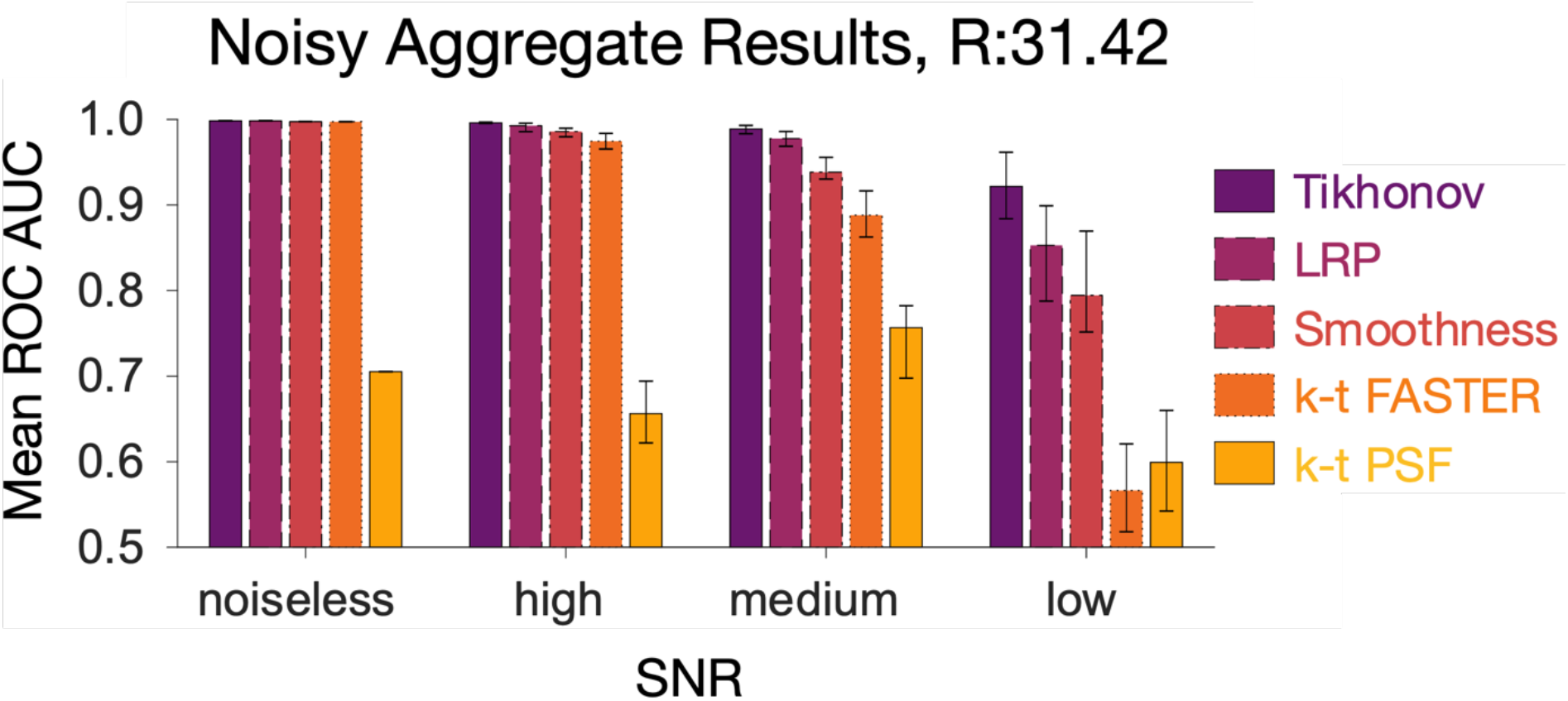
Retrospective dataset B reconstruction AUC results. Each bar represents the mean AUC of five different instantiations of Gaussian noise in k-t space at a specific SNR for a specific reconstruction method, except for the left-hand set, which represent a single noiseless reconstruction. The error bars show the range of AUC values.

**Figure 7:**
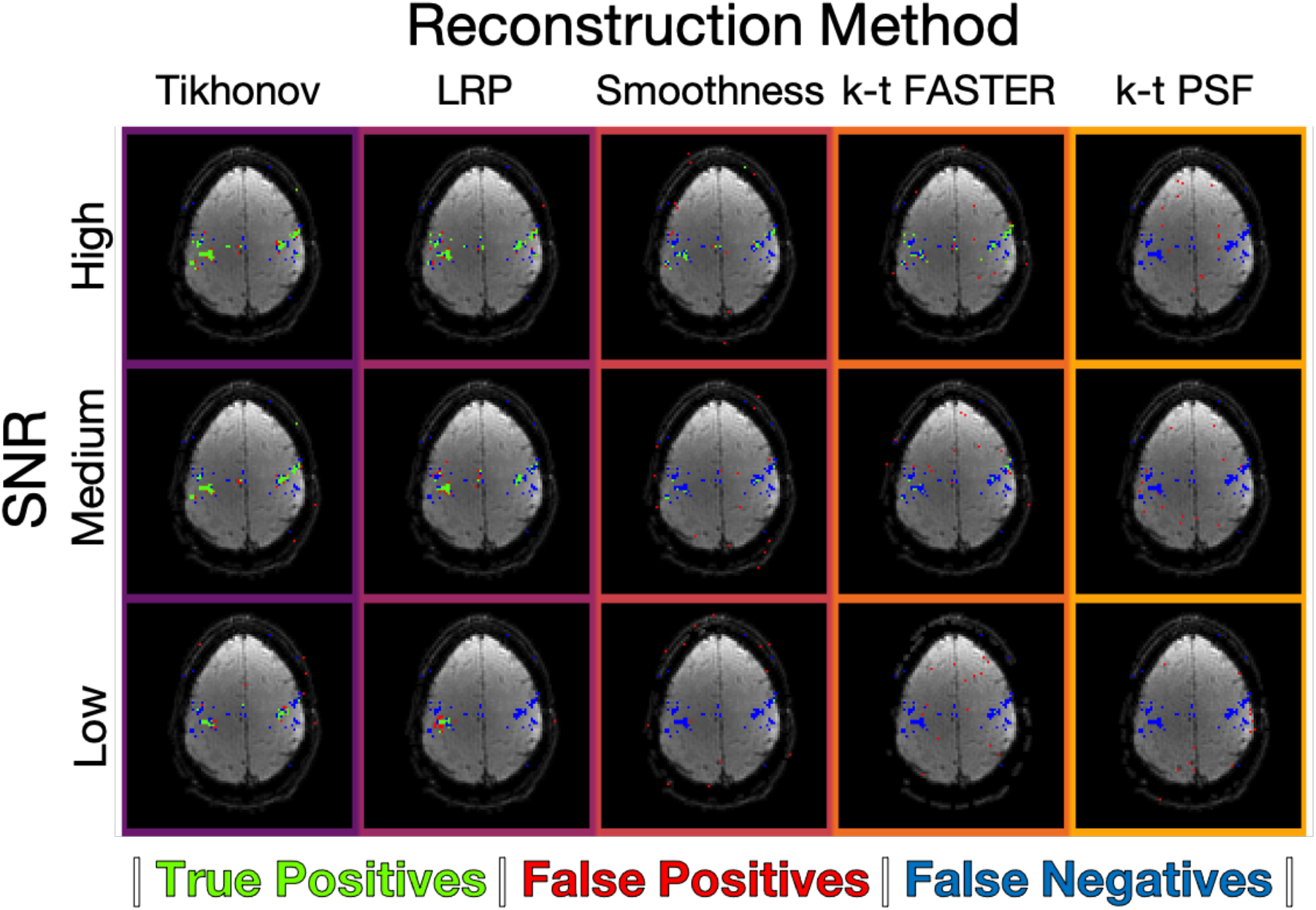
An example activation map at each noise value for each reconstruction method. See Supplementary Figures 3-5 for the full set of activation maps and the individual ROC curves. As with Figures 4-5, green pixels represent true positives, red pixels represent false positives, blue pixels represent false negatives. The z-statistics threshold yielded a false positive rate of 0.15%.

Tikhonov-constrained reconstruction outperformed all other methods, identifying plausible activity even at the lowest SNR tested. LRP and temporal smoothness constraints represent improvements on the previously proposed techniques (k-t FASTER and PSF), with all constrained results better than all k-t FASTER results at medium and low SNR. The k-t FASTER approach appears highly susceptible to noise, with a roughly equivalent noiseless AUC score to the other methods at R=31.42 (figure 5) rapidly decreasing as SNR decreased. The k-t PSF approach failed to capture activation even for the noiseless simulated dataset at this acceleration factor.

### 3.3 Prospective Results

This section presents results on the prospectively under-sampled (“real”) experiments, with three different acceleration factors tested: R=26.2 (6 blades/frame); R=15.7 (10 blades/frame); and R=7.9 (20 blades/frame). The optimal λ values were found to be dependent on both R and the chosen constraint in the prospective dataset (the distribution of reconstruction scores with respect to λ were similar to Figure 3, and so are not shown here). The only exception is that the LRPs were less dependent on λ_T_, with a broader range of values producing scores close to the optimum. Optimal λ values are shown in table 3.

**Table 3:** the optimum λ values in the prospective dataset for each constraint at each acceleration factor. The time in brackets shows the split between the time taken to generate the priors and the final reconstruction. Results with the shortest reconstruction time or within 0.001 of the best ROC AUC score are shown in bold.

| R | *Blades*<br>*Frames* | Method | $\lambda_x$ | $\lambda_T$ | $\lambda_v$ | MEAN RECON TIME<br>(HOURS) | ROC AUC |
| --- | --- | --- | --- | --- | --- | --- | --- |
| **7.85** | 20 | Tikhonov | $10^{-1}$ | $10^{-2}$ | 0 | 2.9 | **0.9915** |
| | | LRP | $10^{-1}$ | $10^{-7}$ | 0 | (1.7+1.6) 3.3 | **0.9913** |
| | | Smoothness | 0 | 0 | $10^{-3}$ | **1.4** | **0.9911** |
|  |  | k-t FASTER | 0 | 0 | 0 | **1.4** | **0.9906** |
| | | k-t PSF | 0 | ∞ | 0 | (1.7+0.3) 2.0 | 0.9884 |
| **15.71** | 10 | Tikhonov | $10^{-1}$ | $10^{-2}$ | 0 | 6.3 | **0.9871** |
| | | LRP | $10^{-1}$ | $10^{-3}$ | 0 | (26.9+6.4) 33.3 | 0.9851 |
| | | Smoothness | 0 | 0 | $10^{+1}$ | 11.2 | **0.9880** |
|  |  | k-t FASTER | 0 | 0 | 0 | **5.8** | 0.9644 |
| | | k-t PSF | 0 | ∞ | 0 | (26.9+0.3) 27.2 | 0.9000 |
| **26.18** | 6 | Tikhonov | $10^{-2}$ | $10^{-1}$ | 0 | **11.6** | 0.9785 |
| | | LRP | $10^{-3}$ | $10^{-7}$ | 0 | (192.3+11.3) 203.6 | 0.9586 |
| | | Smoothness | 0 | 0 | $10^{+2}$ | 29.6 | **0.9875** |
|  |  | k-t FASTER | 0 | 0 | 0 | 13.0 | 0.9410 |
| | | k-t PSF | 0 | ∞ | 0 | (192.3+0.3) 192.6 | 0.4613 |

The ROC curves for the optimal λ at each acceleration factor for each method are shown in Figure 8. The activation maps for every second slice of the R=15.7 and R=26.2 results are shown in Figures 9 and 10 respectively. The full selection of activation maps for all slices and acceleration factors can be seen in Supplementary Figures 6-8.

**Figure 8:**
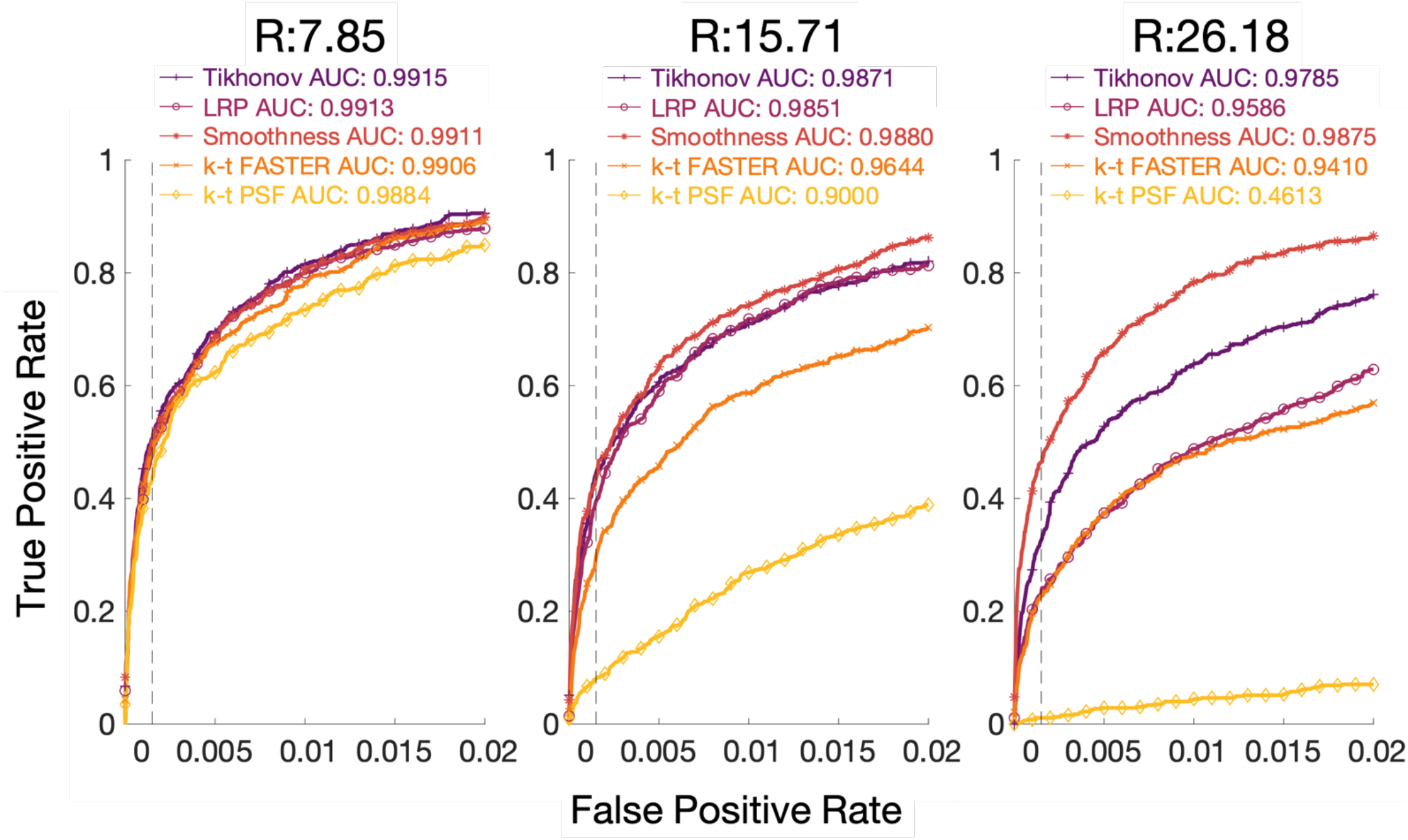
The ROC curves across eight slices for a) R=7.85 (20 blades/frame), b) R=15.71 (10 blades/frame), and c) R=26.18 (6 blades/frame). The ground truth is the long dataset taken under similar experimental conditions, at a threshold of z≥4.8. The false-positive rate is shown on the x-axis up to 0.02, in order to allow visualization of the analytically relevant representation of the activation maps.

**Figure 9:**
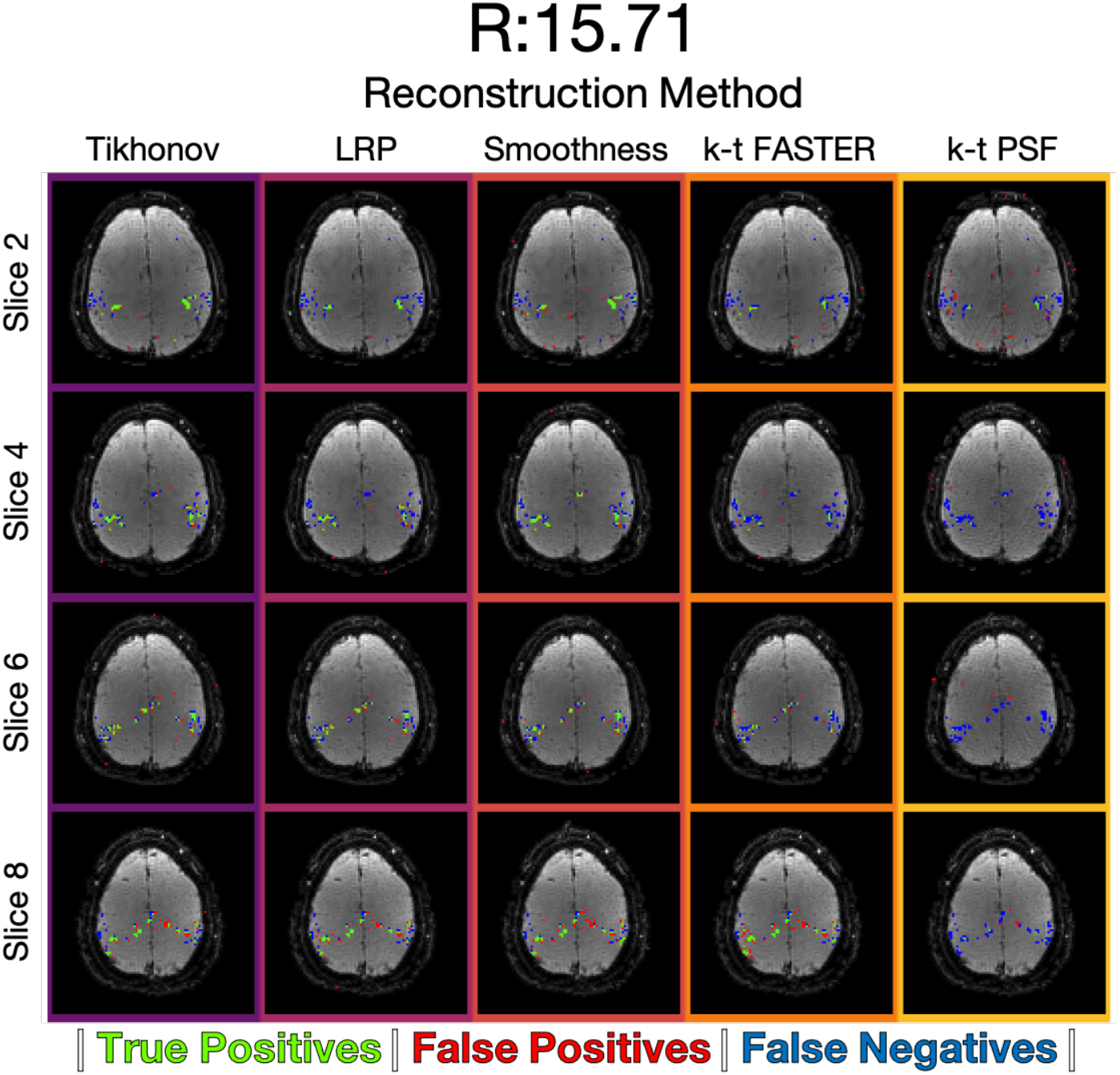
Prospective data, R=15.71. The activation maps for every second slice of the reconstruction, at a threshold defined by a 0.15% volumetric false positive rate. Supplementary Figure 7 shows the activation maps of all slices.

**Figure 10:**
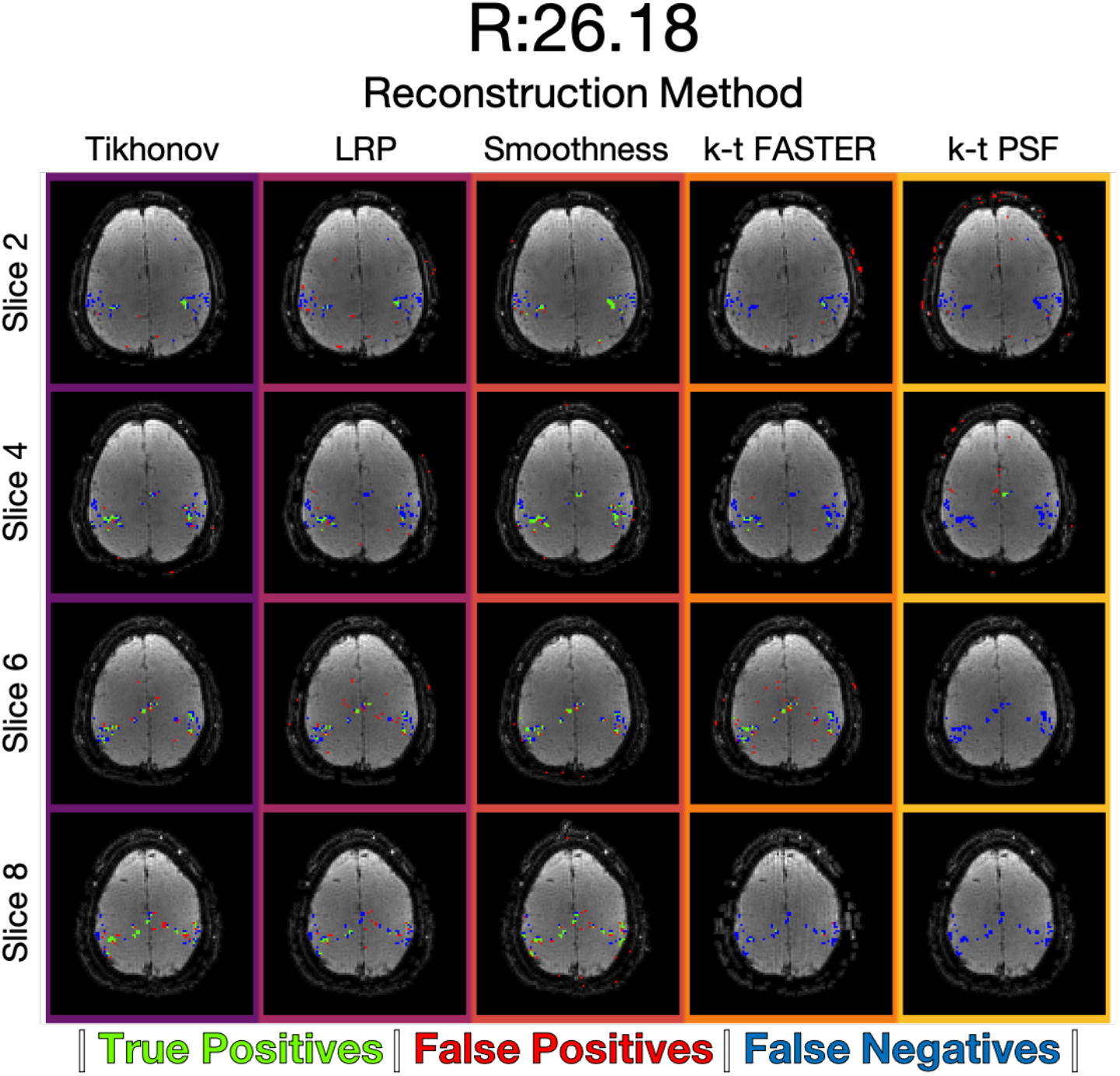
Prospective data, R=26.18. The activation maps for every second slice of the reconstruction, at a threshold defined by a 0.15% volumetric false positive rate. Supplementary Figure 8 shows the activation maps of all slices.

At the lower acceleration factor (R=7.85), all approaches appear approximately equivalent, with k-t PSF performing worst with AUC = 0.9884 and all other methods having AUC > 0.99. At the medium acceleration factors (R=15.71, Figure 9), the soft subspace constraints outperformed k-t FASTER (AUC = 0.9644) and k-t PSF (AUC = 0.90) with AUC > 0.98. At the high acceleration factor (R=26.18, Figure 10), the Tikhonov-constrained results and smoothness results outperformed all other methods with AUCs of 0.9785 and 0.9875 respectively, and the LRP constrained method (AUC = 0.9586) performing similar to k-t FASTER (AUC = 0.9410) at this acceleration factor. Here, the smoothness constraints outperformed the Tikhonov constraints by a score of 0.09, whereas the Tikhonov constraints either performed equivalently or outperformed the smoothness constraints in all other scenarios.

## 4. Discussion

This study demonstrates the impact of three different L2-based constraints in a global low-rank optimization framework for accelerated fMRI data reconstruction. In instances of high acceleration or low SNR, the constrained approaches are able to better identify true regions of activation in a finger-tapping study, as well as producing solutions which more closely map to the spatial and temporal subspaces of a ground truth. These results highlight the viability of non-linear reconstruction frameworks in fMRI that do not rely explicitly on sparse modelling of the BOLD signals.

### 4.1 Comparison between methods

Across the different evaluated datasets a clear trend emerged: the addition of soft subspace-constraints to the k-t FASTER formulation produces improved subspace alignment and ROC AUC scores at high acceleration/low SNR. Collectively, the qualitative and quantitative metrics reveal that very high acceleration factors are possible with these soft constrained-subspace low-rank approaches, in the right conditions. The conditions tested in this paper show that the fMRI signal of interest can be represented by a small number of high-variance components, as elicited with a finger-tapping motor task experiment. The effectiveness of this approach in other, lower-variance examples such as resting-state fMRI or more subtle task fMRI experiments remains to be seen.

The non-linear reconstruction framework only aimed to recover the first 16 components in a low-rank representation of the signal, resulting in feasible reconstructions at very high acceleration due to the reduced matrix degrees of freedom in the estimated output. The acceleration factors reported here (R=26.18 for the prospective dataset) are considerably higher than those reported in previous studies of low-rank fMRI reconstruction using realistic data, which is facilitated largely by the additional soft subspace-constraints. The high acceleration factors in the retrospective dataset A (e.g. R=52.36) were chosen to differentiate between different constraints, and are not considered representative of realistic acceleration factors.

The Tikhonov constraints produced high fidelity reconstructions in both retrospective and prospective under-sampling, even at acceleration factors or SNR levels where other methods began to fail (e.g. the prospective R=26.18/TR=0.3s results, or the low SNR retrospective dataset B results). Additionally, Tikhonov-constrained reconstructions were the fastest to reconstruct out of all the softly constrained reconstructions while its optimal λ pairing could be found through a 1-D parameter search only - reducing the dimensionality of the design constraints.

However, the Tikhonov reconstructions were outperformed by the temporal subspace smoothness approach in the reconstructions of the prospectively under-sampled data, despite that same smoothness approach only providing a relatively small improvement over k-t FASTER in both retrospective datasets. However, the retrospective datasets were constructed under conditions that were favourable for k-t FASTER, without any additional phase modulations or physiological noise (beyond what was in the original dataset). The scale of improvement is also worth noting, with the AUC scores showing Tikhonov outperforming smoothness by an absolute value of +0.3% in the most discriminatory result of retrospective dataset A (R=52.36, 0.9967 vs 0.9938), but smoothness outperforming Tikhonov by +0.9% in the highest acceleration factor tested in the prospective data (R=26.18, 0.9875 vs 0.9785). This smoothness improvement is in addition to the improvement the Tikhonov approach manages over all other methods (+3.75% total over k-t FASTER), while also occurring in the dataset most representative of real data. The outstanding question from these findings is then whether all real-data reconstructions favour smoothing constraints, or are there a set of conditions in real data that would favour Tikhonov constraints?

The low-resolution priors were unable to match the performance of the Tikhonov constraints in any dataset, nor the temporal smoothness in the prospective dataset. The false positives in the LRP-constrained z-stat maps were localized close to the area of interest, indicating the influence of the prior and resulting potential reduction in effective spatial resolution. By comparison, at lower SNR k-t FASTER produced false positives which were less localized to voxels adjacent to true positive activations. As a generalization of the k-t PSF approach, this may reflect the intrinsic limitation of generating priors from low-resolution training data for constraining a high-resolution reconstruction. Furthermore, reconstruction times for the LRP constrained reconstructions were the longest by far.

The k-t PSF method did well at R=7.85 in the real prospective data, and has not to our knowledge been previously tested without sparsity constraints in an fMRI framework. However, the formulation of k-t PSF used in this paper did not produce robust solutions in the other datasets or at the higher acceleration factors tested This is also consistent with the performance of the low-resolution prior method, where both methods that constrained the reconstruction based on a low-spatial resolution temporal basis were not as successful as the other constraints in under-sampled signal recovery.

The optimal regularization factors varied between datasets, and were dependent on SNR for Tikhonov/LRP constraints, and weakly with R. It is clear that a soft constraint can help guide the dataset to improved reconstruction scores, but as with many regularization methods, identification of optimal λ parameters will require some care.

### 4.2 Limitations and Future Work

One limitation of this work is the small sample of datasets used to evaluate the methods, and further testing on additional datasets with physiological noise models or other confounding factors would be needed to establish robustness. This would allow more insight into the robustness of the Tikhonov and smoothness constraints, the optimal λ values, and the impact of coherent noise contamination or auto-regressive noise properties on the different approaches. In addition, further dataset testing could assess the impact of motion. Motion can violate the low-rank assumptions in fMRI, with motion-related variance swamping BOLD fluctuations, and so adequate motion-correction is required. However, a major challenge is that this effect cannot be corrected post-hoc using conventional time-series registration, but needs to correct the k-space data prior to low-rank reconstruction. The data collected for this study was performed on healthy volunteers with very little apparent motion, although the TURBINE k-space trajectory enables motion correction using low spatial resolution navigators [36]. One solution could involve combining TURBINE’s self-navigation capabilities with a joint estimation of the subspaces and motion parameters, leveraging an assumption that a motion-free reconstruction would have the lowest rank or nuclear norm. While the TURBINE acquisition scheme was used to help fulfil the non-uniform sampling density requirement of the LRP constraints, alternative sampling schemes could also be tested to explore how well the smoothness and Tikhonov constraints generalize.

The joint-optimization of two subspaces in alternating minimization provides a flexible reconstruction framework, but could benefit from speeding up. The slowest reconstructions took up to 10s of hours per slice for both Tikhonov and smoothness-constrained reconstruction (Table 2). While Toeplitz Embedding was used to speed up iterative use of the NUFFT [52], [53], the reconstruction code has not been optimized for speed and these computation times could likely be reduced significantly. In addition to code optimization, subproblem parameters such as the convergence factor ε and the number of internal iterations in each linear subproblem (see Supplementary Figure 2) were both chosen to be deliberately conservative for this exploratory analysis and could be fine-tuned for faster reconstructions in future.

## 5. Conclusions

Low-rank reconstructions in fMRI can benefit from additional regularization, particularly at high acceleration factors or in low-SNR regimes. The L2-based constrained-subspace approaches studied here were shown to improve upon methods like k-t FASTER in realistic fMRI data at acceleration factors of R>10, although there is an associated increase in reconstruction time as currently implemented. The improvements with the soft subspace constraints were most apparent at the highest acceleration factor tested (R=26, TR=0.3), and particularly pronounced for the Tikhonov constraints and temporal smoothness constraints.

## Appendix A Pseudocode

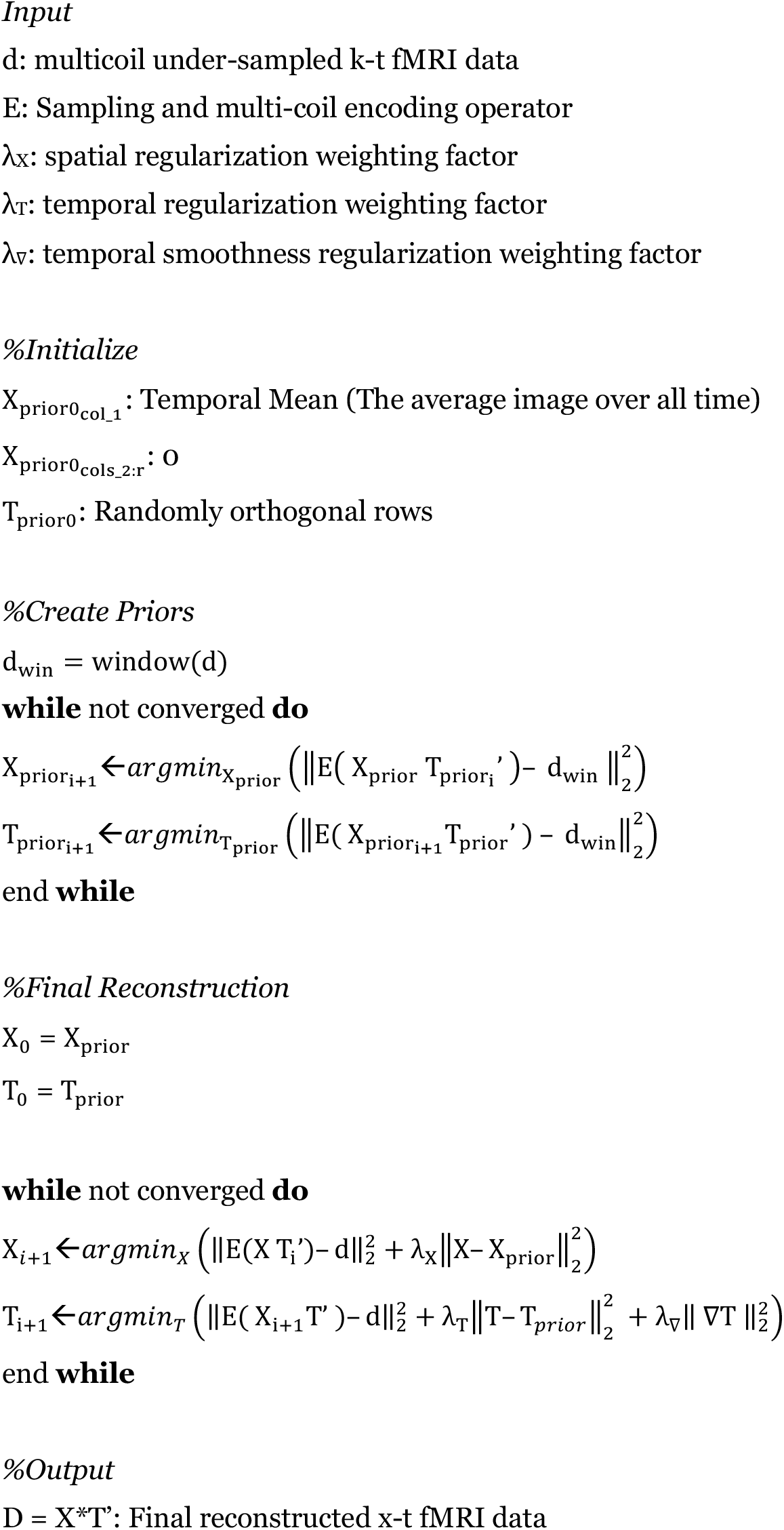

Test code for running the main algorithm of this paper can be found at https://github.com/harrytmason/constrained-lowrank-recon, and the data can be downloaded from ORA once it is made public (it is currently being processed). A link to the data will be provided in the readme file of the code.

## Appendix B Implementation Details

There are a few ways to tackle a k-t space reconstruction problem that constructs a low-rank matrix (e.g. minimizing the nuclear norm: the sum of the singular values [54]; or matrix completion [55]). The approach used in our formulation is known as alternating minimization [38], which reconstructs the decomposed matrices at a fixed rank, pre-selecting an arbitrary low-rank value below the maximum potential rank of the system. Each row in X represents a separate voxel, each row in T represents a frame in time, and the rank is encoded through the columns of both matrices. An additional adaptation employed during prior generation is the forced orthogonalization of the system when alternating between the two subproblems where no alternate regularization exists (e.g. where λ_X_ = λ_T_ = 0).

Our reconstruction problem was solved using the minres.m function in MATLAB R2019a. NUFFT calculations used the Fessler toolbox [37]. Canonical correlations were calculated using the subspacea.m function [48] rather than the inbuilt canoncorr.m function, in order to avoid the extra alignment that occurs during demeaning (which is only significant for low canonical correlation scores).

For windowing, a Tukey parameter of 0.4 was used with full-width half-maximum at 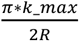. For generation of the priors, a 1D Tukey window was applied along each acquired blade in k-space, and a 2D version of the window was applied to the priors in Cartesian k-t space post prior-generation, but pre-final reconstruction with the full k-space. This ensured no leakage of energy into the higher frequencies, as the windowed data in a consistency term does not strictly enforce the output to only the central k-space.

The overall convergence criterion was a normalized cost function gradient; it was evaluated relative to the cost function at the previous post temporal subproblem iteration after the temporal subproblem in each cycle. The CCS metric was used to establish robustness within a given acceleration factor with respect to the convergence criterion, by reconstructing from different randomly initialized X and T matrices and measuring the agreement of those reconstructions with respect to the principal angles at different levels of convergence. The reconstructions were carried out through a k-t FASTER reconstruction of retrospective dataset A, and are shown in Supplementary Figure 1.

The differing size of the spatial and temporal subproblem means the spatial and temporal problems require different convergence and/or iteration parameters (typically there are 1-2 orders of magnitude more voxels than frames). We chose parameters that made the system spend 10x as long in the spatial subproblem (50 iterations per temporal subproblem, 500 per spatial subproblem, with a subproblem tolerance of 10^−15^ in case of early convergence). The effect of varying the number of iterations of each subproblem against the cycles between the subproblem is shown in Supplementary Figure 2. An internal iteration number of 50 was chosen to guarantee convergence, but this has the potential to be optimized for speed.

Toeplitz embedding exploits the Gram matrix (*E’E*) formed by Fourier encoding to produce a block Toeplitz structure. These can be embedded in block Circulant matrices, which can be fully explained by their first column, and are diagonalized by FFTs. Toeplitz Embedding speeds up the computation from O(N^2^) to O(NlogN). Mark Chiew’s tools for implementing can be found at https://users.fmrib.ox.ac.uk/~mchiew/tools.html.

## Supplementary Figures

**Supplementary Figure 1:**
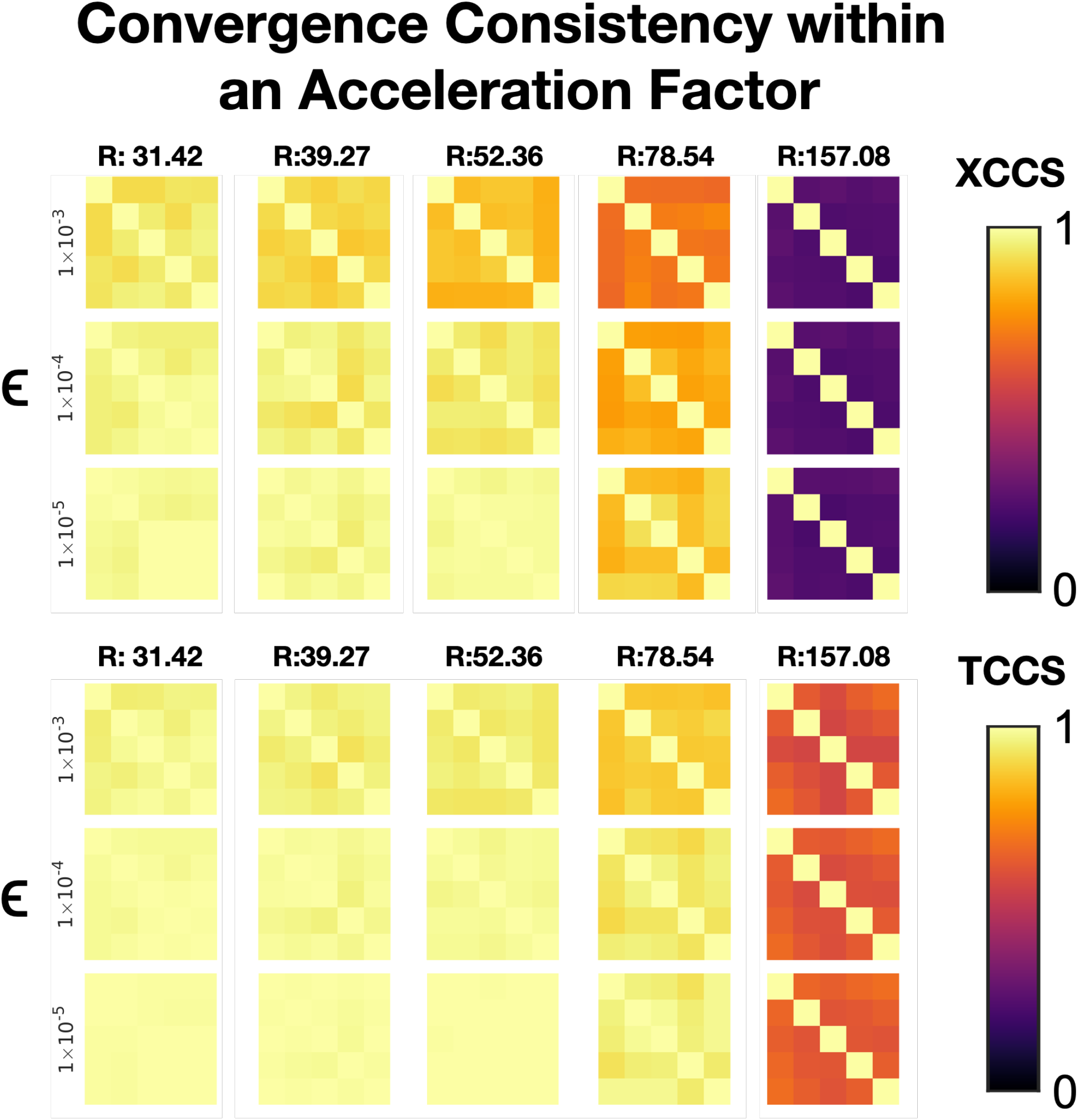
A measure of the robustness of the k-t FASTER algorithm. At each different acceleration factor (columns), five different reconstructions with randomly orthogonal initialization and a temporal mean as the first component were carried out. The result was saved at four different relative absolute gradients of the cost function (rows). The CCS between these different initializations is then shown in each grid, with the diagonal indicating a self-CCS of 1. It is worth noting that random non-orthogonal initializations showed much poorer convergence to a single solution.

**Supplementary Figure 2:**
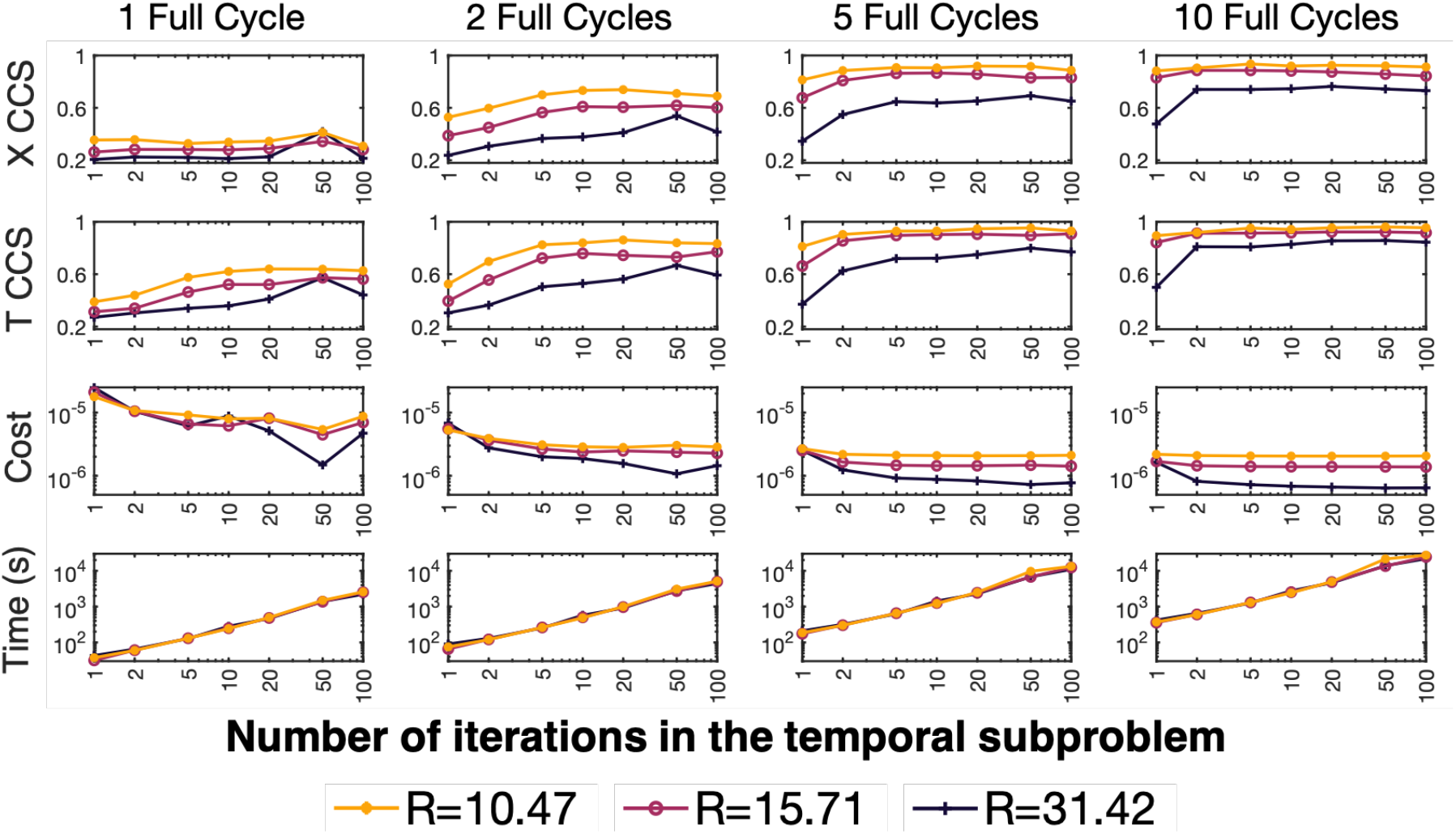
Various criteria (Row 1: X CCS, Row 2: T CCS, Row 3: Cost, Row 4: Time) are used to judge the reconstruction performance with varying iteration numbers in the subproblems. Three different acceleration factors are shown (R=31.42, 5 blades/frame; R=15.71, 10 blades/frame; R=10.47, 15 blades/frame) across a range of cycles (shown in each column). The number of iterations in the spatial subproblem was 10× higher. These results were acquired using alternating minimization k-t FASTER on retrospective dataset A.

**Supplementary figure 3:**
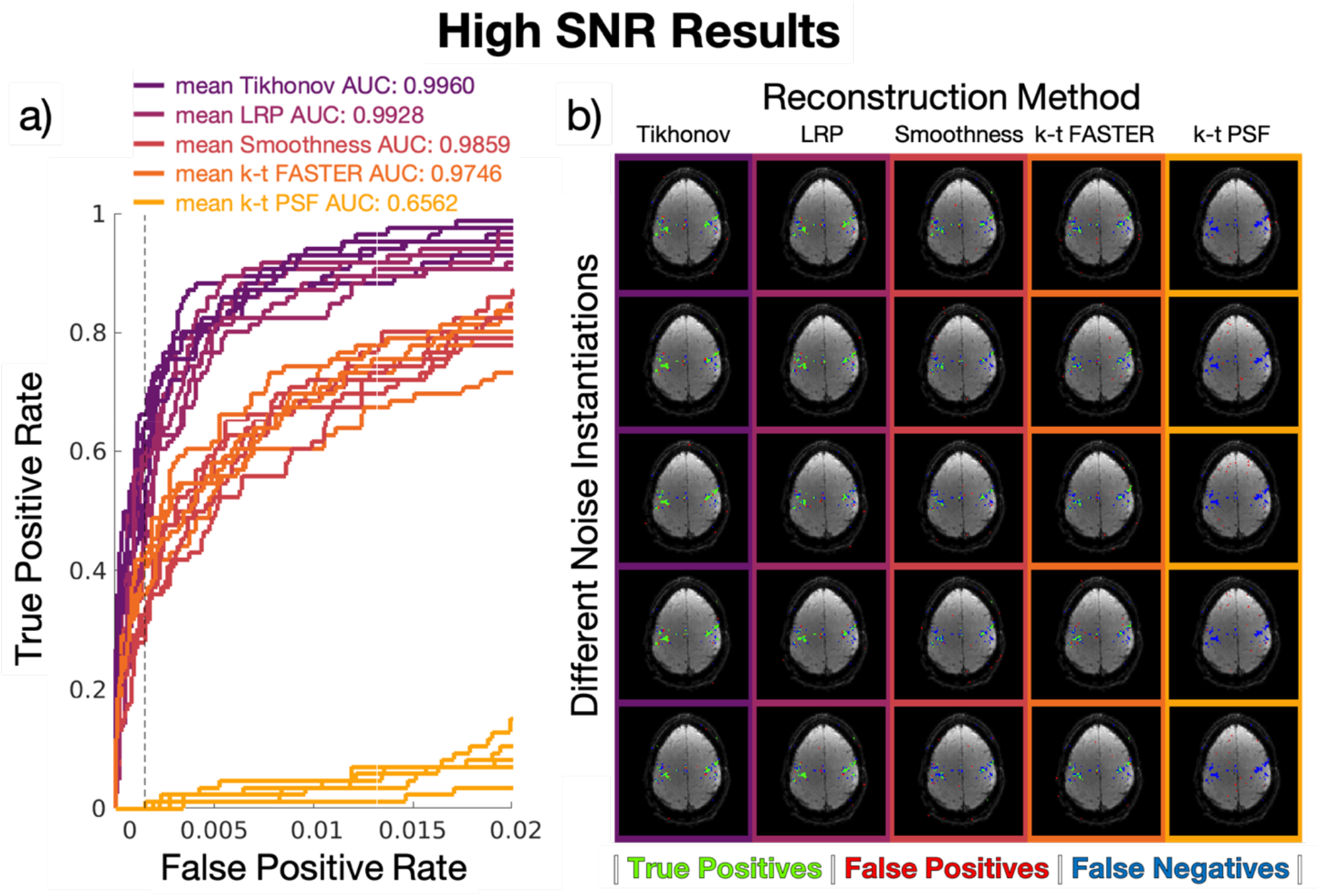
The full set of reconstructions for high SNR in retrospective dataset B. a) The ROC curves for all five instantiations of the noise, when subjected to the different reconstruction methods. The mean AUC across the entire curve is included in the legend. b) The activation maps of all three methods for each individual instantiation of the noise.

**Supplementary figure 4:**
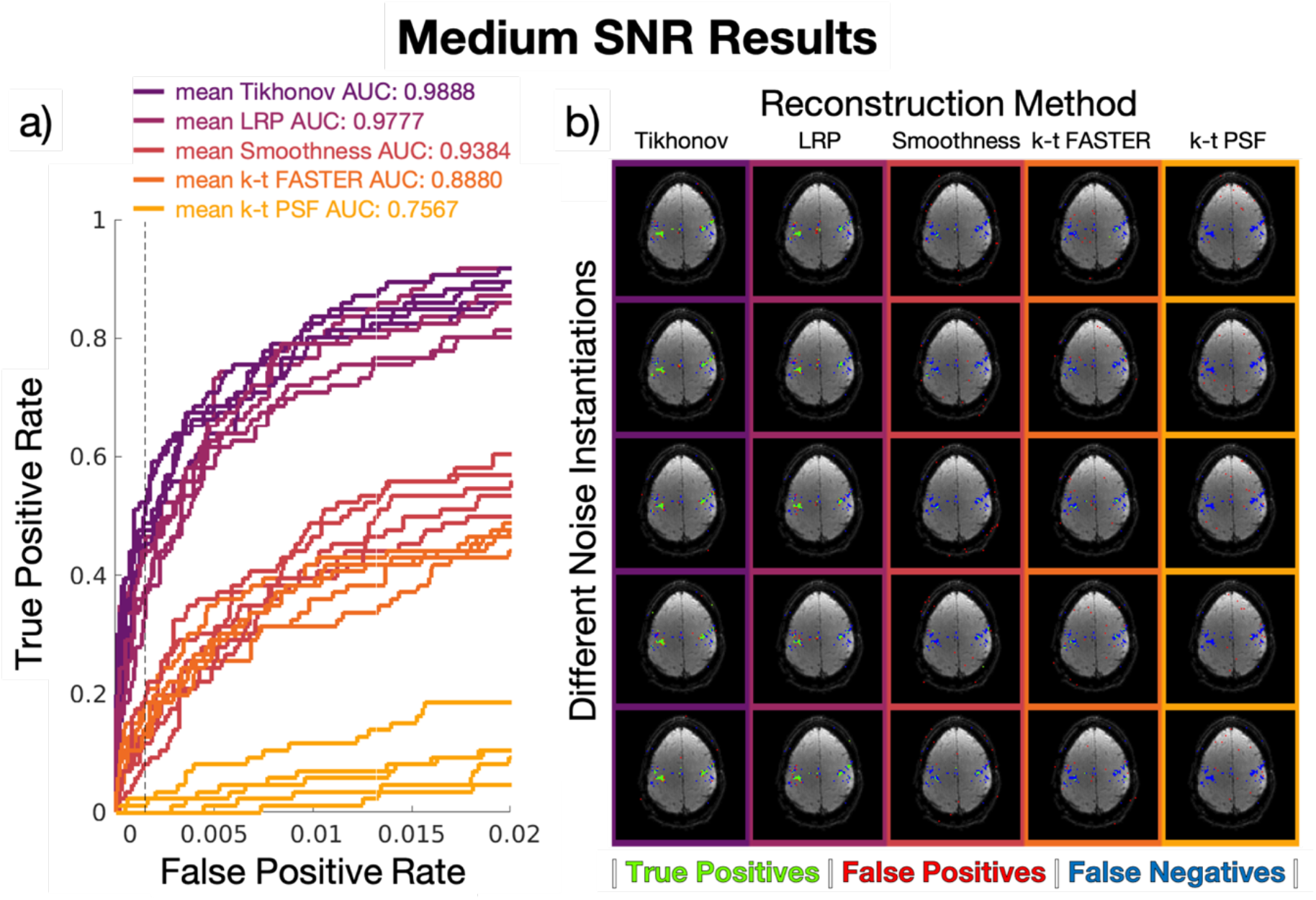
The full set of reconstructions for medium SNR in retrospective dataset B. a) The ROC curves for all five instantiations of the noise, when subjected to the different reconstruction methods. The mean AUC across the entire curve is included in the legend. b) The activation maps of all three methods for each individual instantiation of the noise.

**Supplementary figure 5:**
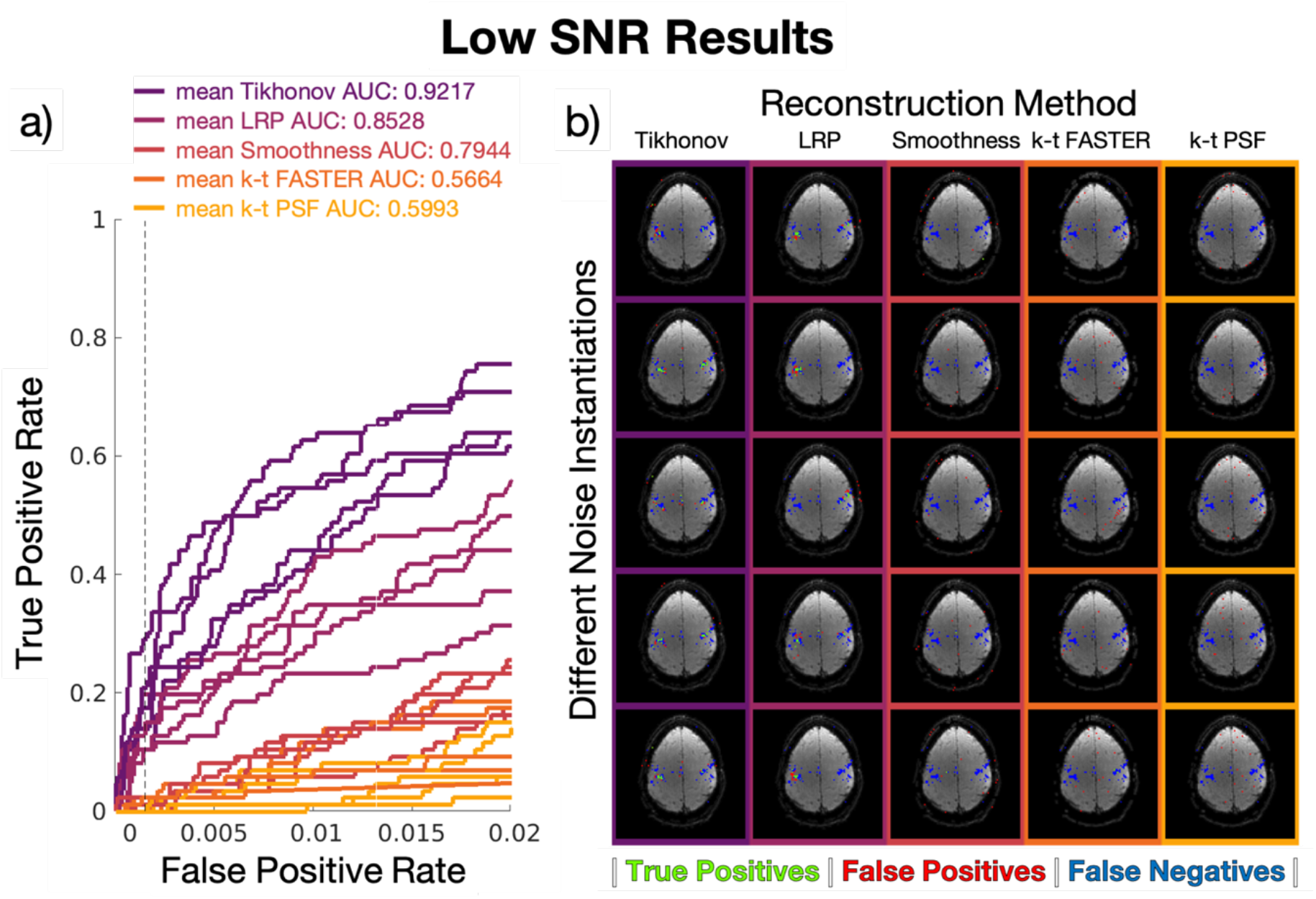
The full set of reconstructions for low SNR in retrospective dataset B. a) The ROC curves for all five instantiations of the noise, when subjected to the different reconstruction methods. The mean AUC across the entire curve is included in the legend. b) The activation maps of all three methods for each individual instantiation of the noise.

**Supplementary figure 6:**
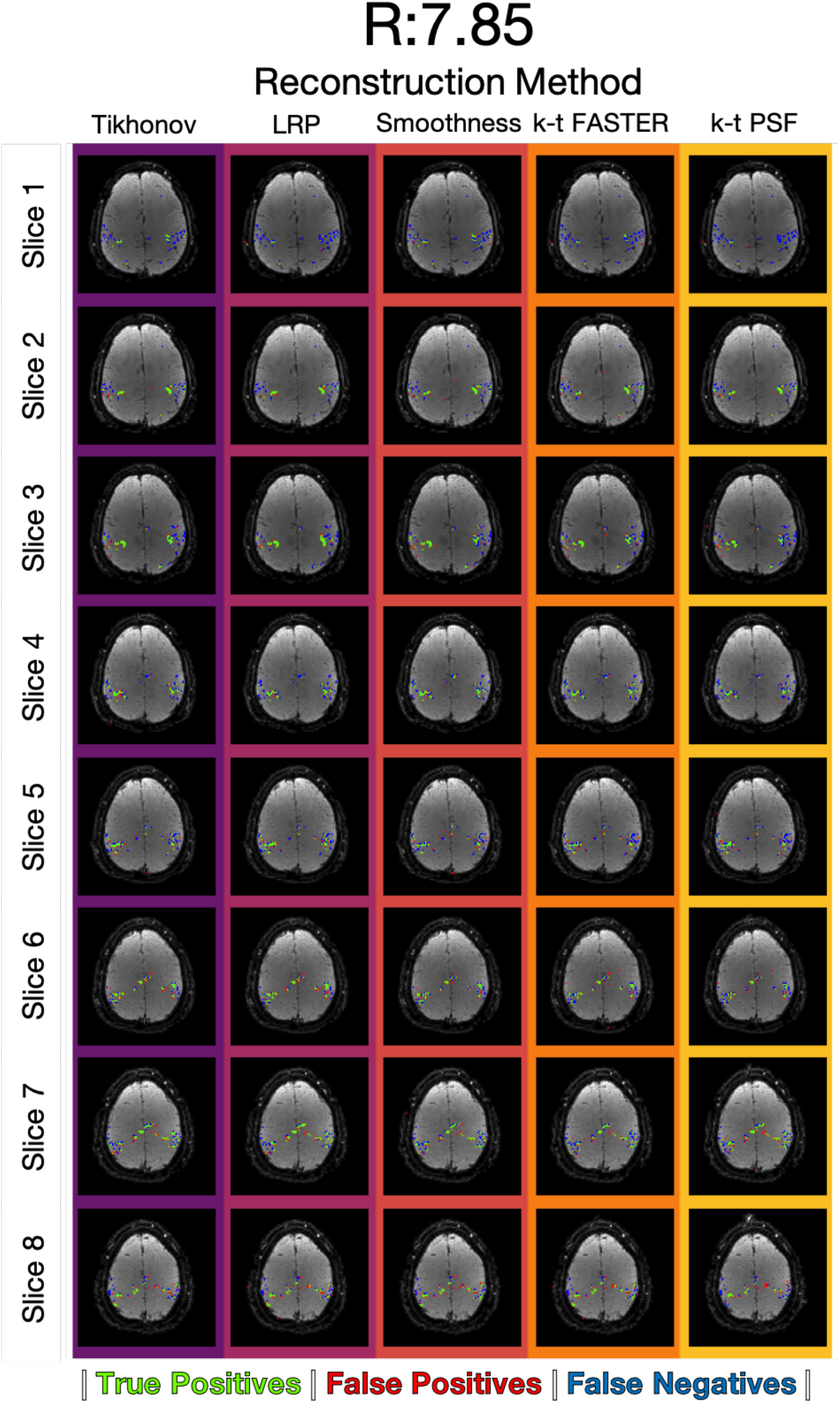
The activation maps for all eight slices for prospective reconstruction at R = 7.85 across the k-t reconstruction methods. The maps were thresholded according to the z-statistic equivalent to a false positive rate of 0.15% (Figure 8). Green pixels represent true positives, blue is false negatives, red is false positives.

**Supplementary figure 7:**
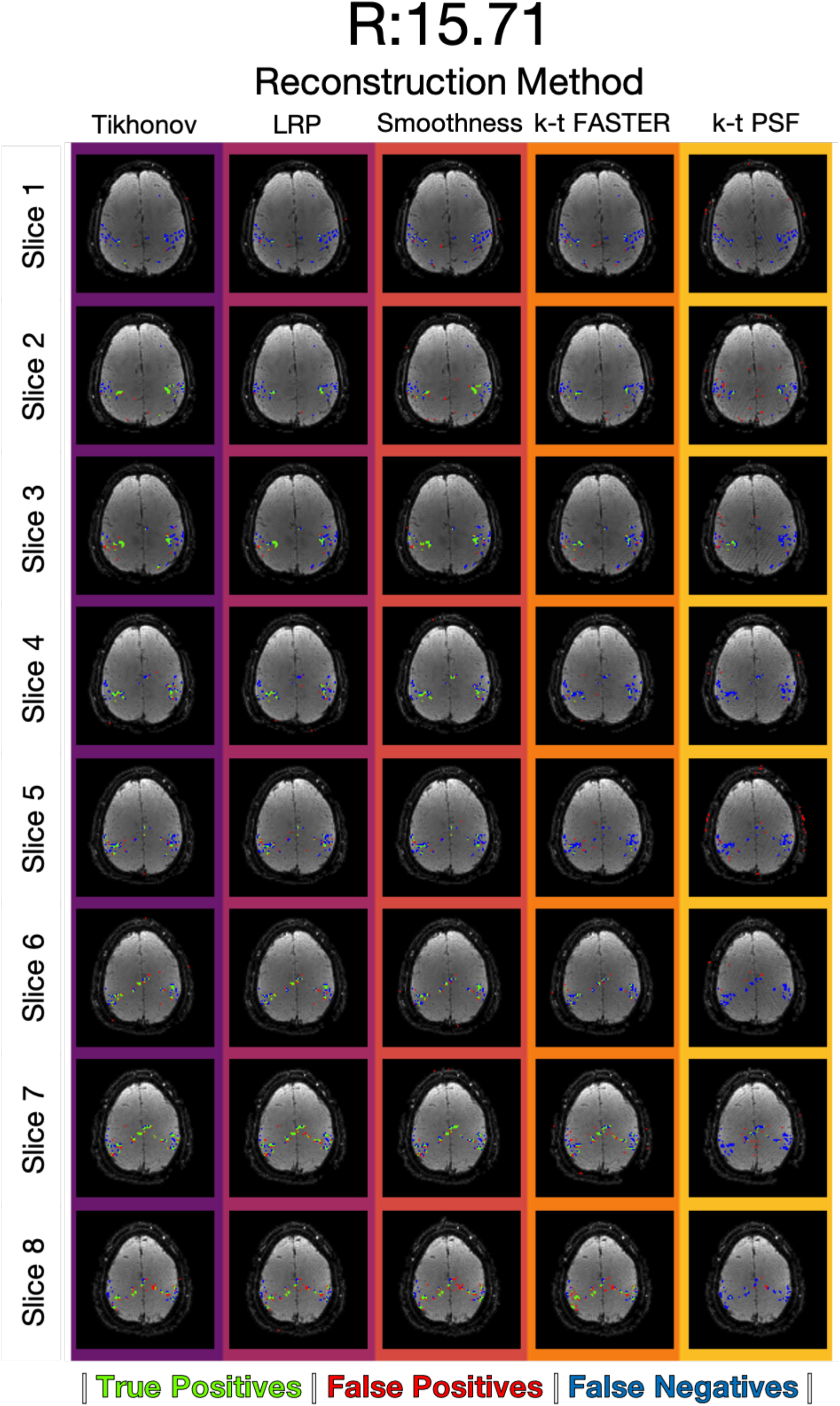
The activation maps for all eight slices for prospective reconstruction at R = 15.71 across the k-t reconstruction methods. The maps were thresholded according to the z-statistic equivalent to a false positive rate of 0.15% (Figure 8). Green pixels represent true positives, blue is false negatives, red is false positives.

**Supplementary figure 8:**
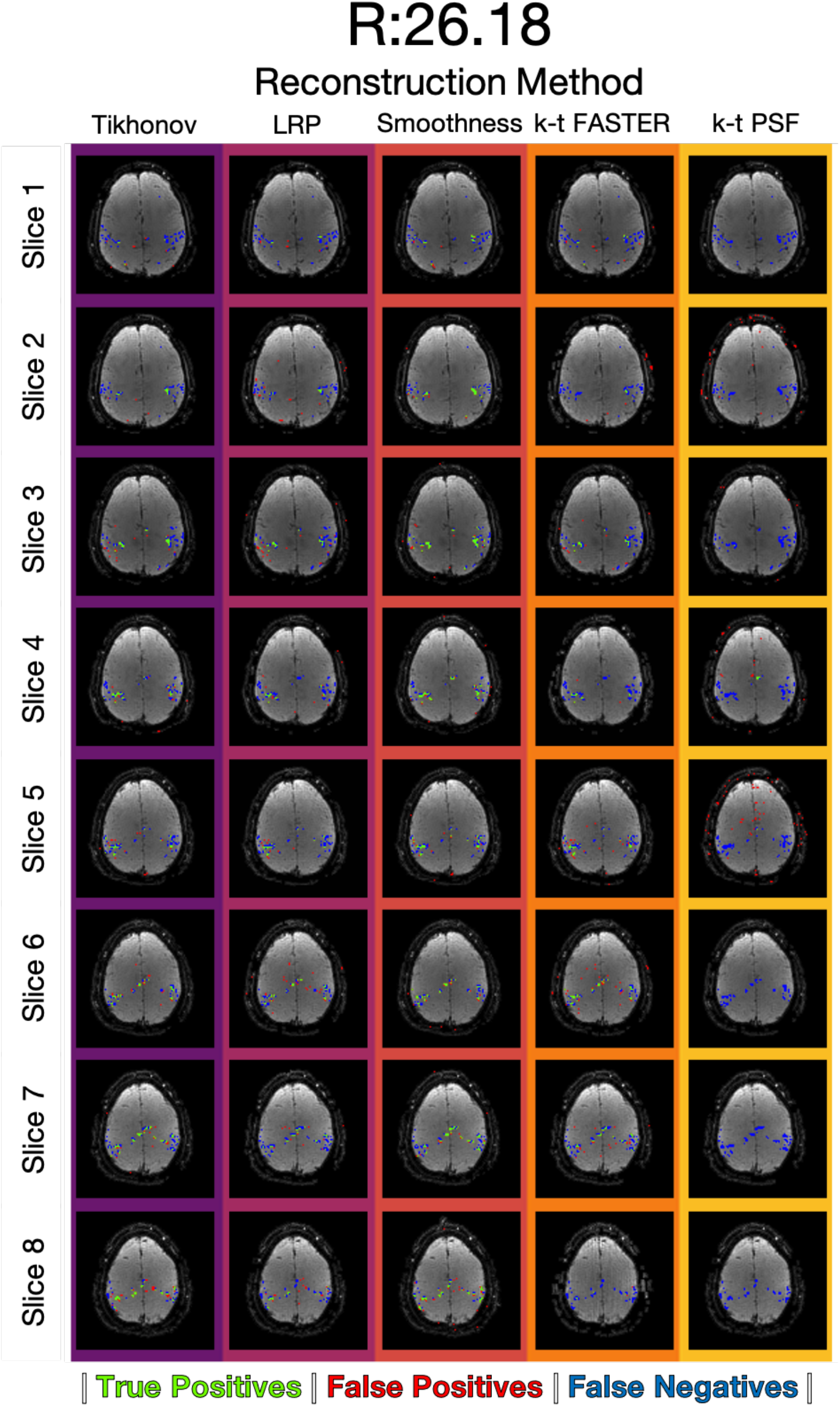
The activation maps for all eight slices for prospective reconstruction at R = 26.18 across the k-t reconstruction methods. The maps were thresholded according to the z-statistic equivalent to a false positive rate of 0.15% (Figure 8). Green pixels represent true positives, blue is false negatives, red is false positives.

## ACKNOWLEDGEMENTS

Computation used the Oxford Biomedical Research Computing (BMRC) facility, a joint development between the Wellcome Centre for Human Genetics and the Big Data Institute supported by Health Data Research UK and the NIHR Oxford Biomedical Research Centre. The views expressed are those of the author(s) and not necessarily those of the NHS, the NIHR or the Department of Health.

This work was supported by funding from the Engineering and Physical Sciences Research Council (EPSRC) and Medical Research Council (MRC) [HTM, grant number EP/L016052/1], the Royal Academy of Engineering (MC, RF201617\16\23) and the Wellcome Trust (KLM, 202788/Z/16/Z). The Wellcome Centre for Integrative Neuroimaging is supported by core funding from the Wellcome Trust (203139/Z/16/Z), and The Wellcome Centre for Human Neuroimaging is supported by core funding from the Wellcome Trust [203147/Z/16/Z].

## Bibliography

[1] K. P. Pruessmann, M. Weiger, M. B. Scheidegger, and P. Boesiger, “SENSE: Sensitivity encoding for fast MRI,” Magn. Reson. Med., vol. 42, no. 5, pp. 952–962, 1999.

[2] M. A. Griswold et al., “Generalized Autocalibrating Partially Parallel Acquisitions (GRAPPA),” Magn. Reson. Med., vol. 47, no. 6, pp. 1202–1210, 2002.

[3] K. Setsompop, B. A. Gagoski, J. R. Polimeni, T. Witzel, V. J. Wedeen, and L. L. Wald, “Blipped-controlled aliasing in parallel imaging for simultaneous multislice echo planar imaging with reduced g-factor penalty,” Magn. Reson. Med., vol. 67, no. 5, pp. 1210–1224, 2012.

[4] M. Barth, F. A. Breuer, P. J. Koopmans, D. G. Norris, and B. A. Poser, “Simultaneous multislice (SMS) imaging techniques,” Magn. Reson. Med., vol. 75, no. 1, pp. 63–81, 2016.

[5] B. Madore, G. H. Glover, and N. J. Pelc, “Unaliasing by Fourier-encoding the overlaps using the temporal dimension (UNFOLD), applied to cardiac imaging and fMRI,” Magn. Reson. Med., vol. 42, no. 5, pp. 813–828, Nov. 1999.

[6] J. Tsao, P. Boesiger, and K. P. Pruessmann, “k-t BLAST and k-t SENSE: Dynamic MRI with high frame rate exploiting spatiotemporal correlations,” Magn. Reson. Med., vol. 50, no. 5, pp. 1031–1042, Nov. 2003.

[7] F. Huang, J. Akao, S. Vijayakumar, G. R. Duensing, and M. Limkeman, “K-t GRAPPA: A k-space implementation for dynamic MRI with high reduction factor,” Magn. Reson. Med., vol. 54, no. 5, pp. 1172–1184, 2005.

[8] S. D. Yun, M. Reske, K. Vahedipour, T. Warbrick, and N. J. Shah, “Parallel imaging acceleration of EPIK for reduced image distortions in fMRI,” Neuroimage, vol. 73, pp. 135–143, 2013.

[9] M. Lustig, D. L. Donoho, and J. M. Pauly, “Sparse MRI: The application of compressed sensing for rapid MR imaging,” Magn. Reson. Med., vol. 58, no. 6, pp. 1182–1195, 2007.

[10] D. J. Holland et al., “Compressed sensing reconstruction improves sensitivity of variable density spiral fMRI,” Magn. Reson. Med., vol. 70, no. 6, pp. 1634–1643, 2013.

[11] O. Jeromin, M. S. Pattichis, and V. D. Calhoun, “Optimal compressed sensing reconstructions of fMRI using 2D deterministic and stochastic sampling geometries,” Biomed. Eng. Online, vol. 11, no. 25, pp. 1–36, 2012.

[12] X. Zong, J. L. Lee, A. Poplawsky, S.-G. Kim, and J. C. Ye, “Compressed Sensing fMRI using Gradient-recalled Echo and EPI Sequences,” Neuroimage, vol. 92, no. 2, pp. 312–321, 2014.

[13] C. Chavarrías, J. F. P. J. Abascal, P. Montesinos, and M. Desco, “Exploitation of temporal redundancy in compressed sensing reconstruction of fMRI studies with a prior-based algorithm (PICCS),” Med. Phys., vol. 42, no. 7, pp. 3814–3821, 2015.

[14] Z.-P. Liang, “Spatiotemporal Imaging with Partially Separable Functions,” IEEE Int. Symp. Biomed. Imaging, vol. 2, pp. 988–991, 2007.

[15] M. Chiew, S. M. Smith, P. J. Koopmans, N. N. Graedel, T. Blumensath, and K. L. Miller, “k-t FASTER: Acceleration of functional MRI data acquisition using low rank constraints,” Magn. Reson. Med., vol. 74, no. 2, pp. 353–364, 2015.

[16] H. Pedersen, S. Kozerke, S. Ringgaard, K. Nehrke, and Y. K. Won, “k-t PCA: Temporally constrained k-t BLAST reconstruction using principal component analysis,” Magn. Reson. Med., vol. 62, no. 3, pp. 706–716, 2009.

[17] H. Jung, K. Sung, K. S. Nayak, E. Y. Kim, and J. C. Ye, “k-t FOCUSS: A general compressed sensing framework for high resolution dynamic MRI,” Magn. Reson. Med., vol. 61, no. 1, pp. 103–116, 2009.

[18] C. Qin et al., “k-t NEXT: Dynamic MR Image Reconstruction Exploiting Spatio-temporal Correlations,” pp. 1–9, 2019.

[19] R. Otazo, E. J. Candès, and D. K. Sodickson, “Low-rank plus sparse matrix decomposition for accelerated dynamic MRI with separation of background and dynamic components,” Magn. Reson. Med., vol. 73, no. 3, pp. 1125–1136, 2015.

[20] A. Y. Petrov, M. Herbst, and V. Andrew Stengerxs, “Improving temporal resolution in fMRI using a 3D spiral acquisition and low rank plus sparse (L+S) reconstruction,” Neuroimage, vol. 157, no. August 2016, pp. 660–674, 2017.

[21] P. Aggarwal and A. Gupta, “Double temporal sparsity based accelerated reconstruction of compressively sensed resting-state fMRI,” Comput. Biol. Med., vol. 91, pp. 255–266, 2017.

[22] Z. Fang, N. Van Le, M. K. Choy, and J. H. Lee, “High spatial resolution compressed sensing (HSPARSE) functional MRI,” Magn. Reson. Med., vol. 76, no. 2, pp. 440–455, 2016.

[23] V. Singh, A. H. Tewfik, and D. B. Ress, “Under-sampled functional MRI using low-rank plus sparse matrix decomposition,” IEEE Int. Conf. Acoust. Speech Signal Process., pp. 897–901, 2015.

[24] P. Aggarwal, P. Shrivastava, T. Kabra, and A. Gupta, “Optshrink LR + S: accelerated fMRI reconstruction using non-convex optimal singular value shrinkage,” Brain Informatics, vol. 4, no. 1, pp. 65–83, 2017.

[25] L. Weizman, K. L. Miller, Y. C. Eldar, and M. Chiew, “PEAR: PEriodic and fixed Rank separation for fast fMRI,” Med. Phys., vol. 44, no. 12, pp. 6166–6182, 2017.

[26] H. Jung, J. C. Ye, and E. Y. Kim, “Improved k-t BLAST and k-t SENSE using FOCUSS,” Phys. Med. Biol., vol. 52, no. 11, pp. 3201–3226, 2007.

[27] M. J. McKeown et al., “Analysis of fMRI data by blind separation into independent spatial components,” Hum. Brain Mapp., vol. 6, no. 3, pp. 160–188, 1998.

[28] A. Hyvärinen, “Fast and Robust Fixed-Point Algorithms for Independent Component Analysis,” IEEE Trans. Neural Networks, vol. 10, pp. 626–634, 1999.

[29] C. F. Beckmann and S. M. Smith, “Probabilistic Independent Component Analysis for Functional Magnetic Resonance Imaging,” IEEE Trans. Med. Imaging, vol. 23, no. 2, pp. 137–152, 2004.

[30] V. Kiviniemi, J. H. Kantola, J. Jauhiainen, A. Hyvärinen, and O. Tervonen, “Independent component analysis of nondeterministic fMRI signal sources,” Neuroimage, vol. 19, no. 2, pp. 253–260, 2003.

[31] G. Salimi-khorshidi, G. Douaud, C. F. Beckmann, M. F. Glasser, L. Griffanti, and S. M. Smith, “Automatic Denoising of Functional MRI Data: Combining Independent Component Analysis and Hierarchical Fusion of Classifiers,” Neuroimage, vol. 44, no. 0, pp. 449–468, Apr. 2015.

[32] F. Lam, B. Zhao, Y. Liu, Z.-P. Liang, M. Weiner, and N. Schuff, “Accelerated fMRI using Low-Rank Model and Sparsity Constraints,” in Proceedings of the International Society for Magnetic Resonance in Medicine 21, 2013, vol. 21, p. 2620.

[33] M. Chiew, N. N. Graedel, J. A. Mcnab, S. M. Smith, and K. L. Miller, “Accelerating functional MRI using fixed-rank approximations and radial-cartesian sampling,” Magn. Reson. Med., vol. 00, pp. 1–12, 2016.

[34] X. Li et al., “Dual-TRACER: High resolution fMRI with constrained evolution reconstruction,” Neuroimage, vol. 164, no. February 2017, pp. 172–182, 2018.

[35] M. Chiew and K. L. Miller, “Improved statistical efficiency of simultaneous multi-slice fMRI by reconstruction with spatially adaptive temporal smoothing,” Neuroimage, vol. 203, no. August, pp. 1–14, Dec. 2019.

[36] N. N. Graedel, J. A. McNab, M. Chiew, and K. L. Miller, “Motion Correction for Functional MRI With Three-Dimensional Hybrid Radial-Cartesian EPI,” Magn. Reson. Med., vol. 78, pp. 527–540, 2017.

[37] J. A. Fessler and B. P. Sutton, “Nonuniform Fast Fourier Transforms Using Min-Max Interpolation,” IEEE Trans. Signal Process., vol. 51, no. 2, pp. 560–574, 2003.

[38] P. Jain, P. Netrapalli, and S. Sanghavi, “Low-rank Matrix Completion using Alternating Minimization,” in Proceedings of the 45th annual ACM symposium on Symposium on theory of computing, 2013, pp. 665–674.

[39] Y. Koren, “Collaborative filtering with temporal dynamics,” Commun. ACM, vol. 53, no. 4, pp. 447–456, 2009.

[40] J. A. McNab, D. Gallichan, and K. L. Miller, “3D steady-state diffusion-weighted imaging with trajectory using radially batched internal navigator echoes (TURBINE),” Magn. Reson. Med., vol. 63, no. 1, pp. 235–242, 2009.

[41] S. Winkelmann, T. Schaeffter, T. Koehler, H. Eggers, and O. Doessel, “An optimal radial profile order based on the golden ratio for time-resolved MRI,” IEEE Trans. Med. Imaging, vol. 26, no. 1, pp. 68–76, 2007.

[42] Y.-C. Kim, S. S. Narayanan, and K. S. Nayak, “Flexible Retrospective Selection of Temporal Resolution in Real-time Speech MRI Using a Golden-Ratio Spiral View Order,” Magn. Reson. Med., vol. 48, no. Suppl 2, pp. 1–6, 2011.

[43] T. Zhang, J. M. Pauly, S. S. Vasanawala, and M. Lustig, “Coil compression for accelerated imaging with Cartesian sampling,” Magn. Reson. Med., vol. 69, no. 2, pp. 571–582, 2013.

[44] M. Buehrer, K. P. Pruessmann, P. Boesiger, and S. Kozerke, “Array compression for MRI with large coil arrays,” Magn. Reson. Med., vol. 57, no. 6, pp. 1131–1139, 2007.

[45] M. Welvaert and Y. Rosseel, “On the definition of signal-to-noise ratio and contrast-to-noise ratio for fMRI data,” PLoS One, vol. 8, no. 11, p. 1010, 2013.

[46] M. Chiew, N. N. Graedel, and K. L. Miller, “Recovering task fMRI signals from highly under-sampled data with low-rank and temporal subspace constraints,” Neuroimage, vol. 174, pp. 97–110, 2018.

[47] Z. Wang, A. C. Bovik, H. R. Sheikh, and E. P. Simoncelli, “Image quality assessment: From error visibility to structural similarity,” IEEE Trans. Image Process., vol. 13, no. 4, pp. 600–612, 2004.

[48] A. V. Knyazev and M. E. Argentati, “Principal Angles Between Subspaces in an A-Based Scalar Product: Algorithms and Estimates,” Soc. Ind. Appl. Math. J. Sci. Comput., vol. 23, no. 6, pp. 2008–2040, 2002.

[49] A. Hyvärinen and E. Oja, “Independent Component Analysis : Algorithms and Applications,” Neural Networks, vol. 13, no. 4–5, pp. 411–430, 2000.

[50] S. M. Smith et al., “Advances in functional and structural MR image analysis and implementation as FSL,” Neuroimage, vol. 23, no. SUPPL. 1, pp. 208–219, 2004.

[51] M. Jenkinson and S. M. Smith, “A global optimisation method for robust affine registration of brain images,” Med. Image Anal., vol. 5, no. 2, pp. 143–156, 2001.

[52] R. Ahmad, C. D. Austin, and L. C. Potter, “Toeplitz embedding for fast iterative regularized imaging,” in Proceedings of the SPIE, 2011, vol. 8051, pp. 1–10.

[53] R. H. Chan and M. K. Ng, “Conjugate Gradient Methods for Toeplitz Systems,” Soc. Ind. Appl. Math., vol. 38, no. 3, pp. 427–482, 1996.

[54] E. J. Candès and B. Recht, “Exact Matrix Completion via Convex Optimization,” Found. Comput. Math., vol. 9, no. 6, pp. 717–772, 2008.

[55] E. J. Candès and T. Tao, “The Power of Convex Relaxation : Near-Optimal Matrix Completion,” IEEE Trans. Inf. Theory, vol. 56, pp. 2053–2080, 2009.

